# Vaccine-induced ICOS+CD38+ cTfh are sensitive biosensors of age-related changes in inflammatory pathways

**DOI:** 10.1101/711911

**Authors:** Ramin Sedaghat Herati, Luisa Victoria Silva, Laura A. Vella, Alexander Muselman, Cecile Alanio, Bertram Bengsch, Raj K. Kurupati, Senthil Kannan, Sasikanth Manne, Andrew V. Kossenkov, David H. Canaday, Susan A. Doyle, Hildegund C.J. Ertl, Kenneth E. Schmader, E. John Wherry

## Abstract

Humoral immune responses are dysregulated with aging but details remain incompletely understood. In particular, little is known about the effects of aging on T follicular helper (Tfh) CD4 cells, the subset that provides critical help to B cells for effective humoral immunity. We previously demonstrated that influenza vaccination increases a circulating Tfh (cTfh) subset that expresses ICOS and CD38, contains influenza-specific memory cells, and is correlated with antibody responses. To directly study the effects of aging on the cTfh response, we performed transcriptional profiling and cellular analysis before and after influenza vaccination in young and elderly adults. Several key differences in cTfh responses were revealed in the elderly. First, whole blood transcriptional profiling defined cross-validated genesets of youth versus aging and these genesets were, compared to other T cells, preferentially enriched in ICOS+CD38+ cTfh from young and elderly subjects, respectively, following vaccination. Second, vaccine-induced ICOS+CD38+ cTfh from the elderly were enriched for transcriptional signatures of inflammation including TNF-NFkB pathway activation. Indeed, we reveal a paradoxical positive effect of TNF signaling on Tfh providing help to B cells linked to survival circuits that may explain detrimental effects of TNF blockade on vaccine responses. Finally, vaccine-induced ICOS+CD38+ cTfh displayed strong enrichment for signatures of underlying age-associated biological changes. Thus, these data reveal key biological changes in cTfh during aging and also demonstrate the sensitivity of vaccine-induced cTfh to underlying changes in host physiology. This latter observation suggests that vaccine-induced cTfh could function as sensitive biosensors of underlying inflammatory and/or overall immune health.

**One sentence summary:** Transcriptional profiling of vaccine-induced circulating T follicular helper cell responding to influenza vaccination reveals age-associated effects on Tfh such as alterations in TNF-NFkB signaling.

## Main Text

Influenza vaccination induces strain-specific neutralizing antibodies in healthy adults, but this process is impaired with aging *(1–4)*. Moreover, immunological aging, that may or may not be directly linked to chronological aging, is also associated with increased morbidity and mortality from infectious diseases *(5, 6)*. Studies of immunological aging and vaccines have identified mechanisms including alterations at the subcellular *(7, 8)*, cellular *(9, 10)*, and systems levels *(11–14)* that lead to altered vaccine responses. Vaccination strategies that can effectively combat age-associated immune dysfunction are of great interest for rational vaccine design. Moreover, identifying strategies to define critical characteristics of overall immune fitness during aging would be of considerable utility for selecting appropriate immunological and other interventions for disease.

The production of class-switched, affinity-matured antibody by B cells is dependent on help from T follicular helper (Tfh) cells in germinal centers (GC) in lymphoid tissues *(15)*, but few studies have evaluated the effects of aging on the GC reaction and GC dependent cellular responses in humans. Suboptimal vaccine responses with aging have been associated with alterations in cellular pathways, such as increased TNF-NFkB signaling or activity *(16)*. For example, several studies observed a weak negative correlation between serum TNF and vaccine responses *(9, 17–19)*. However, therapeutic anti-TNF antibody treatment paradoxically did not result in improved humoral responses and instead led to reduced vaccine-induced antibody responses to pneumococcal and influenza vaccines *(20–22)*. Furthermore, mice deficient in TNF receptor 1 or TNF failed to make GC and were unable to mount sustained IgG responses to immunization *(23–26)*. Thus, the role of TNF signaling in GC and Tfh responses remains poorly understood. Other aspects of Tfh biology that might be affected with aging include impaired cellular activation *(27)*, altered expression of key proteins such as Programmed Death 1 (PD-1, CD279) or Inducible Costimulator (ICOS, CD278) *(10, 27)*, or other pathway-level changes, such as in proliferation or metabolism *(13, 28)*. Understanding these age-associated alterations will be necessary for designing rational vaccines for the elderly.

In humans, directly studying the events leading to productive humoral immunity is challenging because lymphoid tissue is not readily available after vaccination. Several studies have examined a subset of CD4 T cells in human peripheral blood, termed circulating T follicular helper cells (cTfh), that possesses phenotypic, transcriptional, and functional similarities to lymphoid Tfh *(29–34)* and have been used to interrogate vaccine responses. Some features of these cTfh likely reflect events in lymphoid tissue. For example, vaccine-induced changes in cTfh correlate with changes in influenza-specific antibody production *(32–34)*. The subset of cTfh expressing high ICOS and CD38 expands at 7 days after influenza vaccination, correlates with the plasmablast response, and includes recurring influenza-specific T cell receptor clonotypes each year after influenza vaccination *(35)*. Moreover, we have recently demonstrated that these ICOS+CD38+ cTfh can be found in human lymph and share characteristic features of GC Tfh *(36)*. Thus, cTfh may be useful as a cellular biomarker of immunological events including vaccination. Moreover, profiling cTfh may provide opportunities to identify underlying reasons for suboptimal vaccine responses in human populations including the elderly.

Here, we tested the effects of aging on cTfh after influenza vaccination. Consistent with previous studies, the frequency of cTfh subsets responding to vaccination was largely unchanged in older subjects. However, transcriptional profiling revealed key age-associated differences in cTfh responses to influenza vaccination. In particular, transcriptional signatures downstream of TNF-NFkB signaling pathway were enriched in responding ICOS+CD38+ cTfh from elderly compared to young adults at 7 days after vaccination. This transcriptional program was associated with a pro-survival effect in these ICOS+CD38+ cTfh. In addition, transcriptional profiling revealed several pathway-level, age-related differences in activated cTfh responding to vaccination. In particular, on day 7 after vaccination ICOS+CD38+ cTfh from older subjects displayed transcriptional evidence of major alterations in IL2-STAT5 signaling, IL6-STAT3 signaling, and other pathways. Indeed, when examined for general signatures of aging versus youth, ICOS+CD38+ cTfh enriched far better for these signatures than other blood T cell types suggesting that vaccine-induced, or recently activated cTfh are a sensitive biomarker of underlying changes in immune fitness. Thus, these studies identify key changes in cTfh with age that may relate to altered humoral immunity following vaccination and also could provide insight for age-related increases in many autoimmune diseases manifest by autoantibodies. Moreover, activated cTfh may be a useful circulating immune cell type acting as a type of biosensor into underlying changes in immune health and fitness.

## Results

### ICOS+CD38+ cTfh in young and elderly have a transcriptional signature similar to lymphoid Tfh

Antibody responses to vaccination depend on Tfh help in the B cell follicle *(32, 33)*, but the effects of aging on Tfh responses remain poorly understood. We and others have demonstrated that blood cTfh can provide insights Tfh biology relevant to vaccine responses *(32–35)*. Moreover, cTfh expressing ICOS and CD38 expand after influenza vaccination and contain the influenza-specific cells *(32, 35)* and activated Tfh similar to those in the blood can be found in human lymph and have characteristics similar to GC Tfh, suggesting a direct relationship between ICOS+CD38+ cTfh and events in lymphoid tissue *(36)*. In this study, we therefore focused on the ICOS+CD38+ cTfh subset to understand age-related changes in vaccine-induced immunity.

We first sought to investigate the ICOS+CD38+ cTfh subset in young and elderly adults (**Supplemental Table 1**). We defined cTfh as non-naïve CD4+CXCR5+PD-1+ (**Supplemental Figure 1A, Supplemental Table 2**) *(32, 35)*. The frequency of cTfh expressing ICOS and CD38 was similar in young and elderly subjects at baseline and, though the frequency of this cTfh subset increased after influenza vaccination, it was not different between young and elderly subjects (**Figure 1A-C, Supplemental Figure 1B**). The vaccine-induced increase in PD-1 expression was also comparable in both cohorts for the ICOS+CD38+ cTfh population (**Figure 1D, Supplemental Figure 1C-D**). In line with the role of Tfh providing help to B cells, the vaccine-induced fold-change in the ICOS+CD38+ cTfh response correlated with the vaccine-induced fold-change in the plasmablast response, but was not different in young and elderly subjects (young, Pearson r=0.57, P=3.9×10^−3^; elderly, Pearson r=0.67, P=4.8×10^−4^) (**Figure 1E, Supplemental Figure 1E**). Thus, the magnitude of the influenza vaccine induced cTfh responses and their correlation with the plasmablast response was similar in young and elderly subjects.

**Figure 1.**
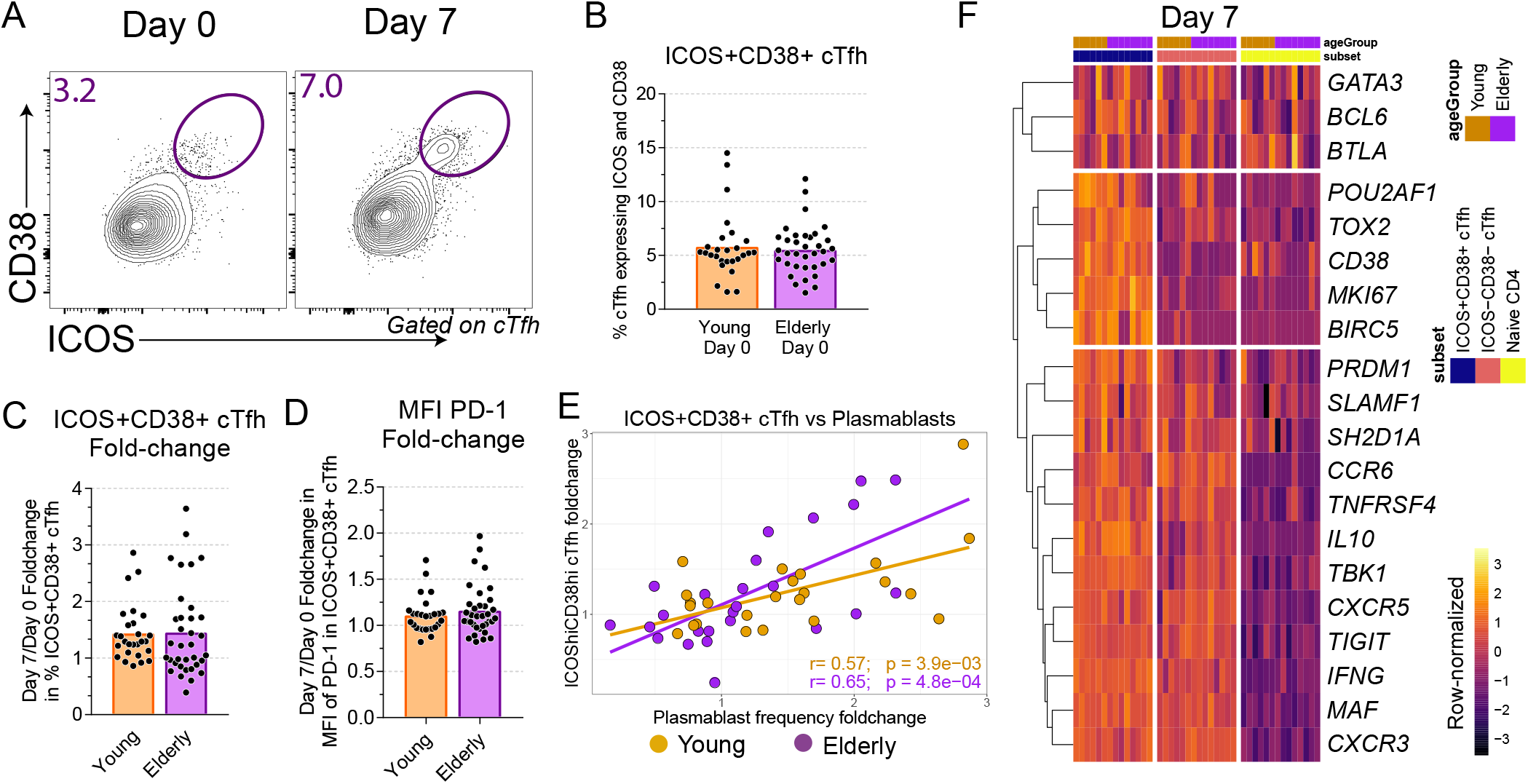
ICOS+CD38+ cTfh express many genes associated with lymphoid Tfh. A cohort of young adults (n=28) and elderly adults (n=35) received influenza vaccination in the Fall of 2014. **A-B.** Flow cytometry (**A**) and summary plots (**B**) shown for days 0 and 7 after vaccination in the frequency of cTfh co-expressing ICOS and CD38 for young (orange, n=27) and elderly (purple, n=35) subjects. **C-D.** Summary plot shown for fold-change between days 0 and 7 for the change in ICOS+CD38+ cTfh frequency (**C**) or geometric MFI of PD-1 (**D**) in ICOS+CD38+ cTfh for young (orange, n=27) and elderly (purple, n=35) subjects. **E.** Correlation shown between the ICOS+CD38+ cTfh frequency fold-change at day 7 compared to day 0, and the plasmablast fold-change at day 7 compared to day 0 for young (orange, r=0.48; P=0.015; Pearson correlation; n=25) and elderly (purple, r=0.54; P=0.005; Pearson correlation; n=26) adults. **F.** Six young and eight elderly adults were randomly selected from the full cohort. PBMC were sorted for ICOS+CD38+ cTfh, ICOS-CD38- cTfh, and naïve CD4 at days 0 and 7 after seasonal influenza vaccination. Log-transformed transcriptional profiling data was queried for selected genes for young (orange bar) and elderly (purple bar) at day 7 after vaccination. Each column represents one unique subject. Heatmap is row-normalized and divided based on hierarchical clustering of genes and CD4 T cell subset.

We next hypothesized that transcriptional profiling may be more sensitive than flow cytometric analysis for revealing differences in cTfh responses. We performed RNAseq on ICOS+CD38+ cTfh, ICOS-CD38- cTfh, and naïve CD4^+^ cells, from young (n=6) and elderly (n=8) adults before and after influenza vaccination (**Supplemental Figure 1F-H**). In both age groups, ICOS+CD38+ cTfh had a distinct transcriptomic signature, with higher expression of genes such as *MAF, BCL6*, and *TIGIT* as compared to naïve CD4, and ICOS+CD38+ cTfh expressed more *CD38, MKI67*, and *POU2AF1* than ICOS-CD38- cTfh (**Figure 1F, Supplemental Figure 1I**). Moreover, the ICOS+CD38+ cTfh displayed greater similarity to lymphoid GC Tfh than the ICOS-CD38- cTfh subset by gene set enrichment analyses (GSEA) *(37)* for human tonsillar GC Tfh *(30)* and mouse lymphoid GC-Tfh *(38, 39)* datasets (**Supplemental Figure 1J-L, Supplemental Table 3**), irrespective of aging. However, there were differences associated with aging at the level of individual gene expression, such as lower expression of *POU2AF1* in elderly at baseline (fold-change 0.76, P=0.058, t-test, 6 young and 8 elderly) (**Figure 1F**) and at day 7 after vaccination (fold-change 0.78, P=0.024, t-test, 6 young and 8 elderly) (**Supplemental Figure 1I**) that suggested that broader age-related differences might be uncovered at the pathway level.

### TNF-NFkB signaling is enriched with aging in ICOS+CD38+ cTfh

To identify age-related differences in the cTfh response to vaccination, we directly compared the transcriptional profiles of ICOS+CD38+ cTfh from young and elderly subjects at day 7 (**Figure 2A**), when the ICOS+CD38+ cTfh population in blood is enriched for influenza-specific cells *(35)*. Differential transcriptional profiles were apparent for ICOS+CD38+ cTfh from young compared to elderly subjects at day 7 (**Supplemental Figure 2A-B**). Differentially expressed genes including *PLCG2, RNF130, RAB2A*, and *ACSL3* that were increased in cTfh from elderly subjects, whereas *CD27, CD6, EIF1AD, EEF2*, and *MAP2K2* were increased in the young (**Supplemental Figure 2B**). CD27, a member of the TNF receptor family, was indeed more strongly expressed at the protein level in the ICOS+CD38+ cTfh from young adults compared to elderly subjects (**Supplemental Figure 2B**). GSEA also identified inflammatory signatures enriched in the ICOS+CD38+ cTfh from the elderly subjects at day 7 post vaccination, with the greatest enrichment for TNF-NFkB signaling (**Supplemental Figure 2C**). Together, these data suggested that the NF-kB signaling pathway was differentially used or engaged by ICOS+CD38+ cTfh in elderly compared to young subjects at day 7 post-vaccination.

**Figure 2.**
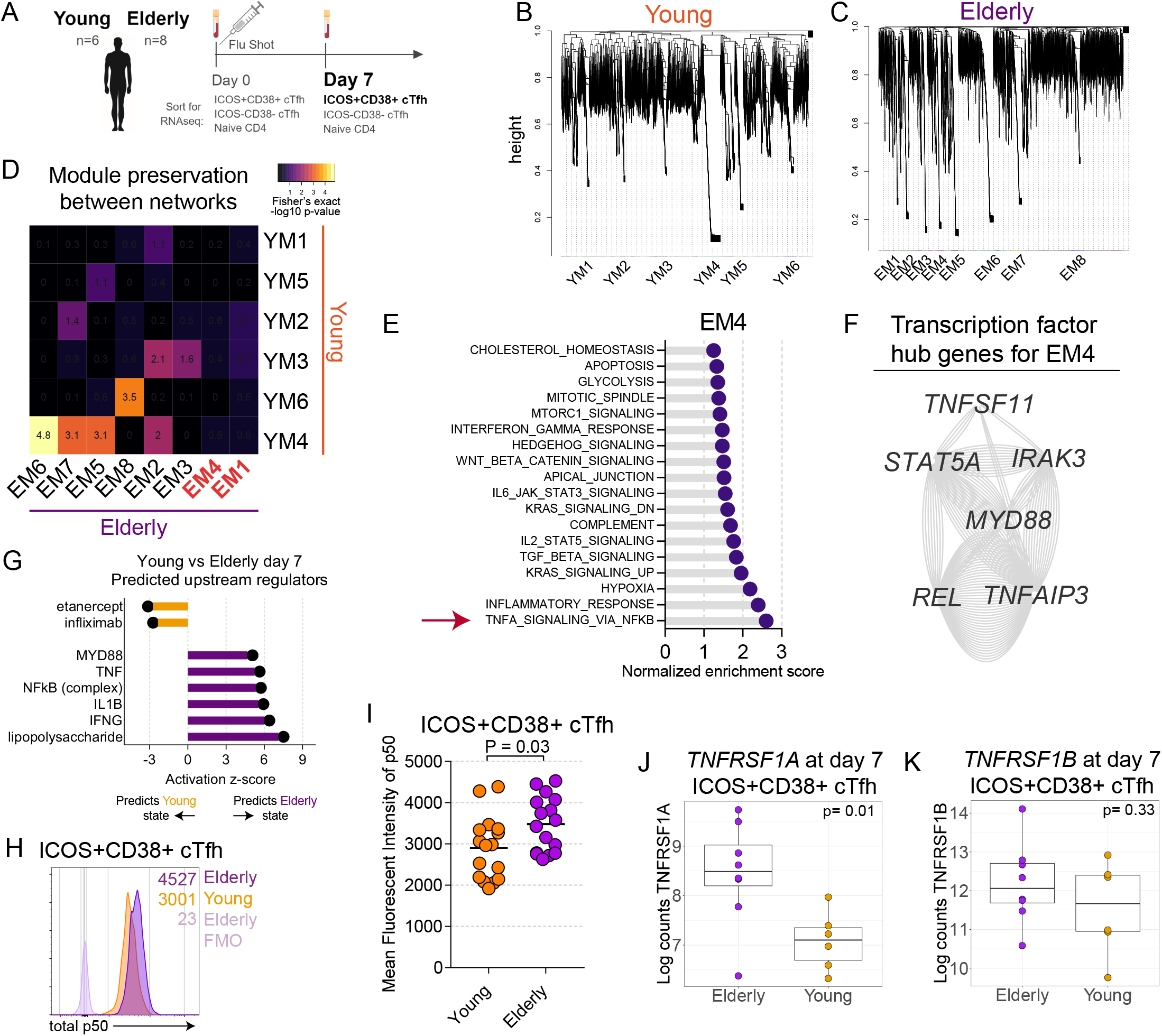
TNF-NFkB pathway is enriched with aging in ICOS+CD38+ cTfh. **A.** Schematic for the RNA-seq analyses indicating comparisons of ICOS+CD38+ cTfh at day 7 with respect to aging, as created with Biorender. **B-C.** Weighted gene correlation network analysis was performed on the ICOS+CD38+ cTfh at day 7 from the six young (**B**) and eight elderly (**C**) subjects. Gene modules are labeled on the dendrogram. **D.** Module preservation analysis was performed using the Fisher’s Exact test. Heatmap color and value indicate the −log_10_(P-value) for the overlap in genes in modules. **E.** Genes were ranked based on module membership for Elderly modules EM1 and EM4, then GSEA analysis was performed using the MSigDB HALLMARK collection. Positive enrichment scores indicate enrichment for genes in module EM4. **F.** Genes with module membership >0.80 for module EM4 were filtered for transcription factors and selected transcription factors displayed as a multiple association network using GeneMANIA without gene prediction. **G.** Predicted upstream regulators were analyzed for the differential expression comparison of ICOS+CD38+ cTfh at day 7 for young and elderly subjects using Ingenuity Pathway Analysis. Activation z-scores are shown for the indicated terms. Positive activation z-scores predict the elderly state whereas negative scores predict the young state. **H.** Example flow cytometry plot from one young and one elderly subject for total NFkB p50 protein. Numbers on the plot indicate geometric MFI for p50. **I.** Total NFkB p50 was measured by flow cytometry for the ICOS+CD38+ cTfh at baseline in young (orange) and elderly (purple) adults (P=0.031; t-test; n=16 in each group). **J-K.** Gene expression for *TNFRSF1A* (**J**, P=0.01, t-test, n=6 for young; n=8 for elderly) and *TNFRSF1B* (**K**, P=0.33, t-test, n=6 for young; n=8 for elderly) shown at day 7 from log_2_-transformed counts data for young (orange) and elderly (purple).

These observations provoked the hypothesis that inflammatory signals were driving transcriptional network differences in the vaccine-induced cTfh from elderly subjects. To investigate this possibility, we next applied weighted gene correlation network analysis (WGCNA) *(40)* to more deeply interrogate underlying transcriptional networks of vaccine induced cTfh from young versus elderly subjects. At day 7 after vaccination, six transcriptional modules were identified for the ICOS+CD38+ cTfh subset isolated from young adults, whereas eight transcriptional modules were identified from the elderly (**Figure 2B-C, Supplemental Table 4**). Although most modules in young overlapped with one or more modules in the elderly, two modules in the elderly (EM1 and EM4) did not have significant overlap with any module in the young adults (**Figure 2D, Supplemental Figures 2D-E**). To evaluate the underlying biology of these modules, we used GSEA on genes ranked by module membership for EM1 and EM4 (**Figure 2E, Supplemental Figure 2F**) and found TNF-NFkB signaling to be the highest-scoring pathway associated with EM4. After filtering the EM4 gene list for transcription factors *(41)*, we identified a dense network of six hub genes, *IRAK3, MYD88, TNFAIP3, STAT5A, REL* and *TNFSF11*, that had direct relevance to the NF-kB signaling pathway (**Figure 2F, Supplemental Table 5**). Additionally, Ingenuity Pathway analysis (IPA) for upstream regulators predicted LPS, IL1B, NFkB (complex), TNF, and MYD88 to be more activated in the elderly compared to young adults in ICOS+CD38+ cTfh at day 7 following vaccination (**Figure 2G, Supplemental Figure 2G**). In addition, several other inflammatory pathways were strongly enriched in EM4 including Inflammatory Response, TGF-β signaling, IL-2 STAT5 signaling, and IL-6 JAK STAT3 signaling as were signatures of hypoxia and complement (**Figure 2E**). Module EM1 also enriched for inflammatory pathways as well as type I and type II interferon pathways (**Supplemental Figure 2F**). Together, these results identified coordinated transcriptional changes reflecting inflammatory signaling in the elderly.

We then used gene set variation analysis (GSVA) *(42)* to investigate whether the plasma TNF concentrations had any relationship with the increase of the TNF-NFkB geneset signature in elderly patients 7 days after immunization (**Supplemental Figure 2H**). However, no correlation was observed between the GSVA score for the TNF-NFkB gene set and the serum TNF concentration (**Supplemental Figure 2I**), suggesting that serum TNF alone was not likely to have induced the observed TNF-NFkB transcriptional signature.

We next tested if there was evidence for constitutive upregulation and/or activation of the NFkB pathway *ex vivo*. Indeed, NFKB1 (p50) protein expression was elevated in ICOS+CD38+ cTfh from elderly compared to young adults (**Figure 2H-I, Supplemental Figure 2J**) suggesting a greater activation of, or ability to activate, this pathway in response to inflammatory mediators in the ICOS+CD38+ cTfh from aged individuals. TNF signaling occurs via interaction of soluble or membrane-bound TNF with TNF receptors, CD120a and CD120b (TNFR1 and TNFR2, respectively), with soluble TNF signaling more strongly through TNFR1 than TNFR2 *(43)*. To test whether cTfh differentially expressed TNFR1 or TNFR2, transcriptional profiling data for CD4 T cell subsets was evaluated for TNFR1 (gene name *TNFRSF1A*) and TNFR2 (gene name *TNFRSF1B*). Indeed, ICOS+CD38+ cTfh had higher gene expression for TNFR1 and TNFR2 than other CD4 subsets at baseline (**Supplemental Figure 2K**). Following vaccination, although expression of *TNFRSF1B* was similar between young and elderly subjects, *TNFRSF1A* expression in ICOS+CD38+ cTfh was increased in elderly compared in young adults (**Figure 2J-K, Supplemental Figure 2L**). Thus, several components of the TNF signaling pathway are elevated in vaccine induced ICOS+CD38+ cTfh from elderly compared to young adults consistent with an increase in transcriptional signatures downstream of this signaling pathway in elderly compared to young adults.

### TNF signaling promotes Tfh-B cell interactions

TNF-NFkB signaling with aging has been associated with age-related immune dysfunction *(44)*. Here, our transcriptional and network analyses highlighted a role for increased NF-kB signaling with aging within ICOS+CD38+ cTfh at day 7 postvaccination. We hypothesized that this TNF-NFkB signaling may adversely affect Tfh-B cell interactions. To test this idea we used a cTfh-B cell coculture system *(32)*. Supernatants from these co-cultures revealed a correlation between subject age and TNF in the supernatant after 7 days of co-culture (**Figure 3A**, Pearson r=0.35, P=0.048). Because these co-cultures used the same source of naïve allogeneic B cells from a single young donor, these data indicate greater production of TNF that occurred during *in vitro* cTfh-B cell interactions was due to cTfh from the elderly subjects. Together, these data indicated an age-associated upregulation of the NFkB signaling pathway in the ICOS+CD38+ cTfh after vaccination that was associated with increased TNF production during Tfh-B cell interactions in the elderly.

**Figure 3.**
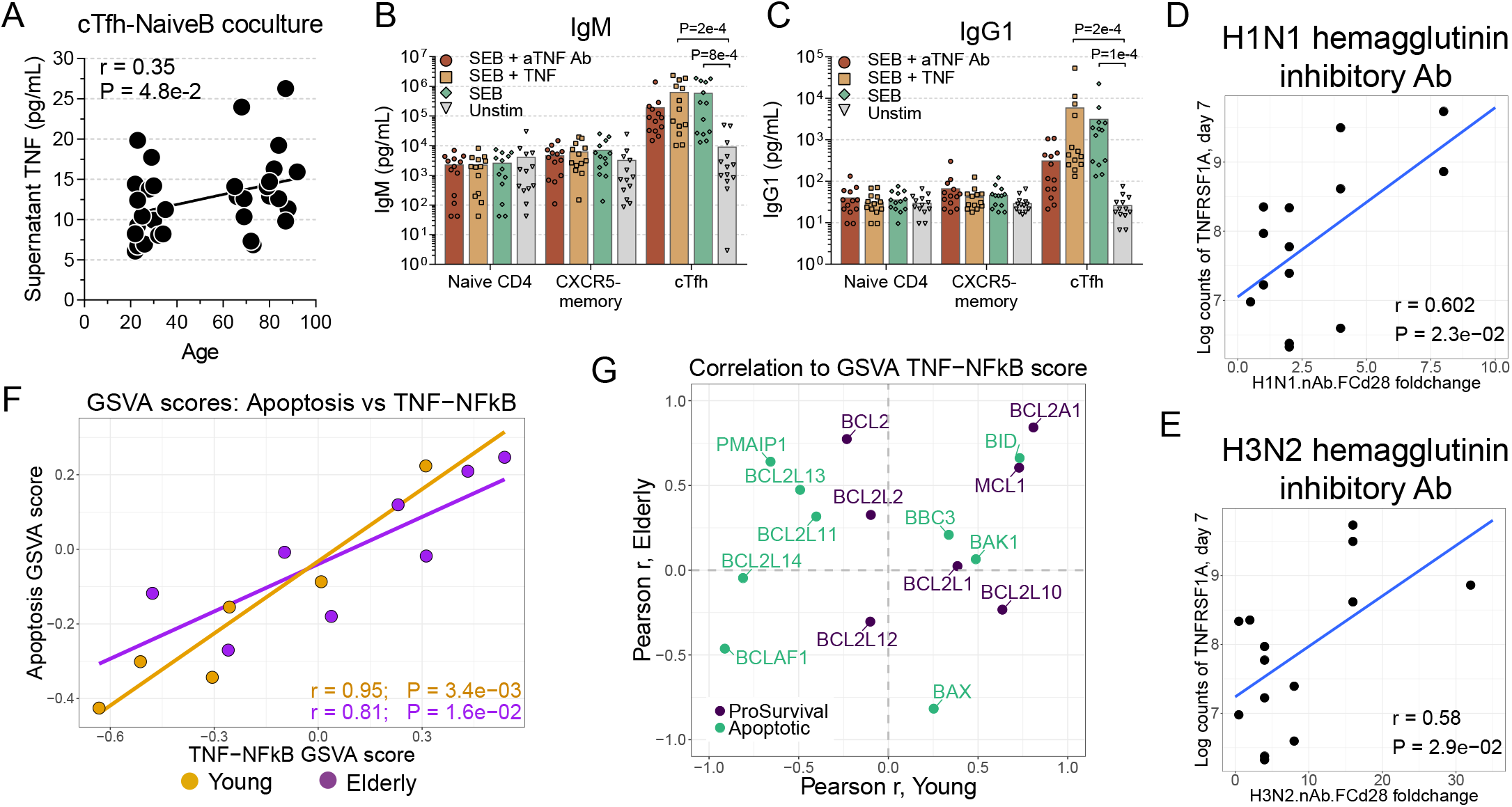
TNF signaling promotes Tfh-B cell interactions. **A.** Supernatant TNF was measured in co-culture of total cTfh from adults with allogeneic naïve B cells from one common young adult after 7 days of stimulation with 0.1 μg/mL Staphylococcal enterotoxin B (P=0.048; Pearson r=0.35; n=33). **B-C.** PBMC from young adults were freshly isolated and sorted for co-culture with autologous naïve B cells (CD3^−^ CD19^+^CD27^lo^IgD^+^). Supernatant IgM (**B**) and IgG1 (**C**) were measured after 7 days, as shown for the following conditions: unstimulated (grey), SEB alone (0.5 μg/mL, green), SEB with recombinant human TNF (125 ng/mL, tan), or SEB with αTNF antibodies (2 μg/mL, sienna) (one-way ANOVA with Freidman’s test; n=12 per group). **D-E.** Correlation shown for the *TNFRSF1A* in ICOS+CD38+ cTfh at day 7 compared with the H1N1-specific (**D**) HAI titer (p=0.029; Pearson r=0.58; n=14) or H3N2-specific (**E**) HAI titer (p=0.023; Pearson r=0.60; n=14). **F.** Correlation of GSVA scores for the Apoptosis and NF-kB genesets for ICOS+CD38+ cTfh at day 7 for young (orange, P=3.4×10^−3^, Pearson r=0.95, n=6) and elderly (purple, P=1.6×10^−2^, Pearson r=0.81, n=8). **G.** Pearson correlation coefficients for BCL family member genes compared to GSVA scores for TNF-NFkB signaling in ICOS+CD38+ cTfh at day 7 for young (x-axis) and elderly (y-axis) adults. Genes were classified as either pro-survival (dark blue) or apoptotic (sea green).

To evaluate the functional role of TNF signaling in the Tfh-B cell interaction, these *in vitro* B cell help assays were extended as previously described *(29, 30, 32)*, using autologous naïve B cells and CD4 T cells from young adults. As expected cTfh were the major supplier of B cell help, as evidenced by higher IgM and IgG1 in supernatant after SEB stimulation compared to controls (**Figure 3B-C**). Soluble, but not membrane-associated TNF, is thought to be needed for proper GC formation *(45)*. To further test the impact of TNF on the ability of cTfh to provide help, supplemental recombinant TNF protein or α-TNF antibody was added at the beginning of the coculture. Adding TNF did not augment B cell help and antibody production compared to SEB alone (**Figure 3B-C**). In contrast, addition of α-TNF blocking antibodies reduced supernatant IgG1 concentrations 10-fold compared to SEB alone (**Figure 3C**). The differentiation of naïve B cells into CD19^+^CD38^+^CD20^lo^ plasmablasts was also reduced in the presence of α-TNF antibodies, but addition of TNF had no effect (**Supplemental Figure 3A**). IgG1 production by naïve B cells stimulated *in vitro* with α-IgM and soluble trimeric CD40L did not change upon addition of α-TNF antibodies (**Supplemental Figure 3B**), suggesting that reduced antibody production by B cells in the context of blockade of TNF signaling in the presence of cTfh was through effects of TNF signaling on Tfh. Indeed, live cell counts for T and B cells decreased when α-TNF antibodies were added to the co-cultures (**Supplemental Figure 3C-D**) or when the TNF converting enzyme inhibitor TAPI-O *(46)* was used (**Supplemental Figure 3E**), suggesting a potential pro-survival role for TNF signaling. Overall, these data suggested a role for TNF-NFkB signaling in promoting cellular survival in the context of Tfh-B cell interactions.

Given the co-culture data, we hypothesized that TNF-NFkB signaling may be associated with a beneficial effect on the humoral response *in vivo*. Influenza vaccination was associated with higher expression of *TNFRSF1A* in ICOS+CD38+ cTfh in the elderly compared to young adults following vaccination (**Figure 2J-K**). Thus, we next compared expression of *TNFRSF1A* and *TNFRSF1B* to the neutralizing antibody responses. Examining young and elderly subjects together, *TNFRSF1A* expression by ICOS+CD38+ cTfh (but not other CD4 subsets) positively correlated with the fold-change in the strain-specific neutralizing antibody titers at day 28 compared to day 0 (**Figure 3D-E**). These data support the notion of a positive role for TNF-NFkB signaling in Tfh-B cell interactions. Moreover, these observations are consistent with observations that clinical TNF blockade resulted in reduced vaccine immunogenicity *(20, 21)*.

Since TNF blockade was associated with reduced survival of cTfh and B cells in cocultures *in vitro*, we next asked if there was an association between TNF signaling and cell death *in vivo*. Indeed, GSVA scores for TNF-NFkB signaling correlated strongly with apoptosis, similarly for both age groups (**Figure 3F**). This positive correlation seemed paradoxical given the *in vitro* data above. However, cells did not appear to be undergoing active regulated cell death *(47)*, as the cellular membrane of ICOS+CD38+ cTfh was intact based on exclusion of viability dye (**Supplemental Figure 1A**), and ICOS+CD38+ cTfh of young and elderly adults did not demonstrate reduced mitochondrial membrane potential compared to other CD4 T cell subsets (**Supplemental Figure 3F**). Resistance to apoptosis could be linked to the induction of pro-survival genes. Indeed, expression of anti-apoptotic genes *BCL2A1* and *MCL1* correlated with the GSVA scores for TNF-NFkB signaling in all subjects (**Figure 3G, Supplemental Figure 3G**). Similarly, expression of survivin (gene name *BIRC5*, **Figure 1F, Supplemental Figure 3H**), a cIAP family member that can restrain caspase-3 activation, was high in ICOS+CD38+ cTfh. Thus, together with the reduced viability *in vitro* with TNF blockade these data suggest that TNF-NFkB signaling may be necessary to counterbalance pro-apoptotic effects and foster cell survival circuits.

### Vaccination elicits discordant transcriptional pathway responses with aging in ICOS+CD38+ cTfh

To better understand the full spectrum of transcriptional changes in cTfh during aging, we analyzed the vaccine-induced transcriptional signatures in ICOS+CD38+ cTfh on day 7 post vaccination compared to day 0 for young versus elderly subjects. Overall, influenza vaccination was associated with relatively modest changes in transcriptional profiles in ICOS+CD38+ cTfh from young and elderly (**Figure 4A**) suggesting that prevaccination ICOS+CD38+ cTfh share many transcriptional characteristics with these cells post vaccination.

**Figure 4.**
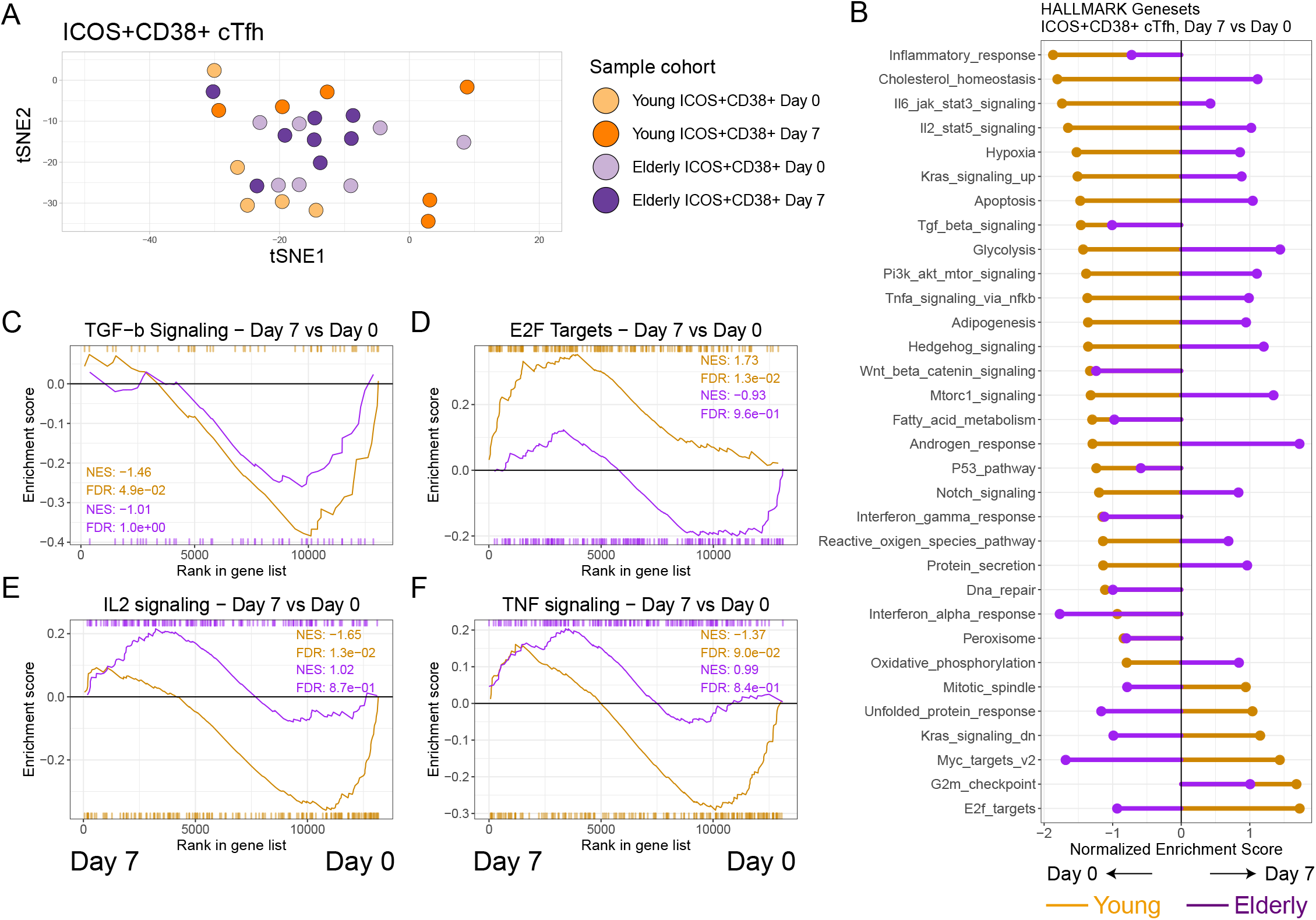
Age-dependent differences in transcriptional profiles of ICOS+CD38+ cTfh. **A.** tSNE analysis was performed for all CD4 T cell subsets in the full dataset. ICOS+CD38+ cTfh are shown before and after vaccination for young (orange shades) and elderly (purple shades). **B.** Aggregated GSEA results are shown for the selected Hallmark genesets for the comparison of day 7 versus day 0 for ICOS+CD38+ cTfh in young (orange) and elderly (purple). Positive enrichment scores indicate enrichment at day 7. **C-F.** GSEA analysis shown for the comparison of day 7 versus day 0 for the ICOS+CD38+ cTfh subset from young (orange) and elderly (purple) for the MSigDB Hallmark collection, for TGFβ signaling (**C**), E2F targets (**D**), IL-2 signaling (**E**), and TNF-NFkB signaling (**F**) genesets.

We then used pathway enrichment combined with pre-ranked GSEA to compare transcriptional signatures from ICOS+CD38+ cTfh from young versus elderly. Clear enrichment patterns were observed for many key gene sets. However, overall the enrichment patterns were only concordant between young and elderly subjects for ~1/3 of the Hallmark gene sets examined (**Figure 4B, Supplemental Figure 4A, Supplemental Table 3**) suggesting distinct biology for this cell type in elderly versus young subjects. For example, TGFβ signaling, displayed reduced enrichment at day 7 compared to day 0 in ICOS+CD38+ cTfh from both young and elderly adults (**Figure 4C**). In contrast, in young adults there was positive enrichment (associated with upregulation at day 7) for the E2F targets geneset in ICOS+CD38+ cTfh, whereas elderly had negative enrichment for this pathway (**Figure 4D**). The opposite pattern was observed for IL-2-STAT5 (**Figure 4E**) and TNF-NFkB signaling (**Figure 4F**), both of which were enriched in the elderly compared to the young. In general, ICOS+CD38+ cTfh from the young enriched positively in signatures of regulation of proliferation and Myc biology, but negatively for signatures of inflammatory signaling. In contrast, ICOS+CD38+ cTfh from the elderly enriched positively for metabolic pathways and inflammatory signaling. These results demonstrate multiple pathway-level differences with aging in ICOS+CD38+ cTfh after influenza vaccination and specifically highlight a potential role for inflammatory pathways.

### Age-related differences preferentially revealed by CD4 T cell subset analysis

We next hypothesized that the age-related differences in cellular pathways may be distinct in different CD4 T cell subsets. We therefore compared enrichment for nine Hallmark genesets by GSEA in naïve CD4 T cells, ICOS-CD38- cTfh and ICOS+CD38+ cTfh from young and elderly subjects before (d0) and after (d7) influenza vaccination (**Figure 5A-I, Supplemental Figure 5A-B, Supplemental Table 3**). These analyses revealed several key findings. First, consistent with our data in ICOS+CD38+ cTfh (**Figure 1**), most pathways showed minimal differences with aging at baseline, a finding pronounced for naïve CD4 T cells. Two exceptions included a moderate enrichment of the Myc targets biological pathway (**Figure 5D**) in ICOS-CD38- cTfh and ICOS+CD38+ cTfh from the elderly at baseline and an enrichment of oxidative phosphorylation (**Figure 5F**) in the ICOS-CD38- cTfh from the young subjects at day 0. Second, the overall similarity at baseline was contrasted by substantial differences following influenza vaccination. Of all genesets, 57% changed in enrichment direction (i.e. from young to elderly or *vice versa*) for ICOS+CD38+ cTfh between days 0 and 7. However, one of the most clear patterns from these analyses was the uncovering of an underlying bias for inflammatory signatures including TNF-NFkB, Inflammatory Response and IL-2-STAT5 gene sets to be preferentially enriched in CD4 T cells of the elderly subjects, often with greatest enrichment in the ICOS+CD38+ cTfh subset (**Figure 5A-C, J**). Third, ICOS+CD38+ cTfh showed the greatest dynamic range out of the three T cell subsets, based on the difference with aging in the day 7 and day 0 normalized enrichment scores per subset (**Figure 5J**). Whether these differences reflect differential systemic induction of inflammation upon vaccination, distinct abilities of T cells from elderly versus young subject to induce transcriptional responses related to these pathways, or both, remains to be determined.

**Figure 5.**
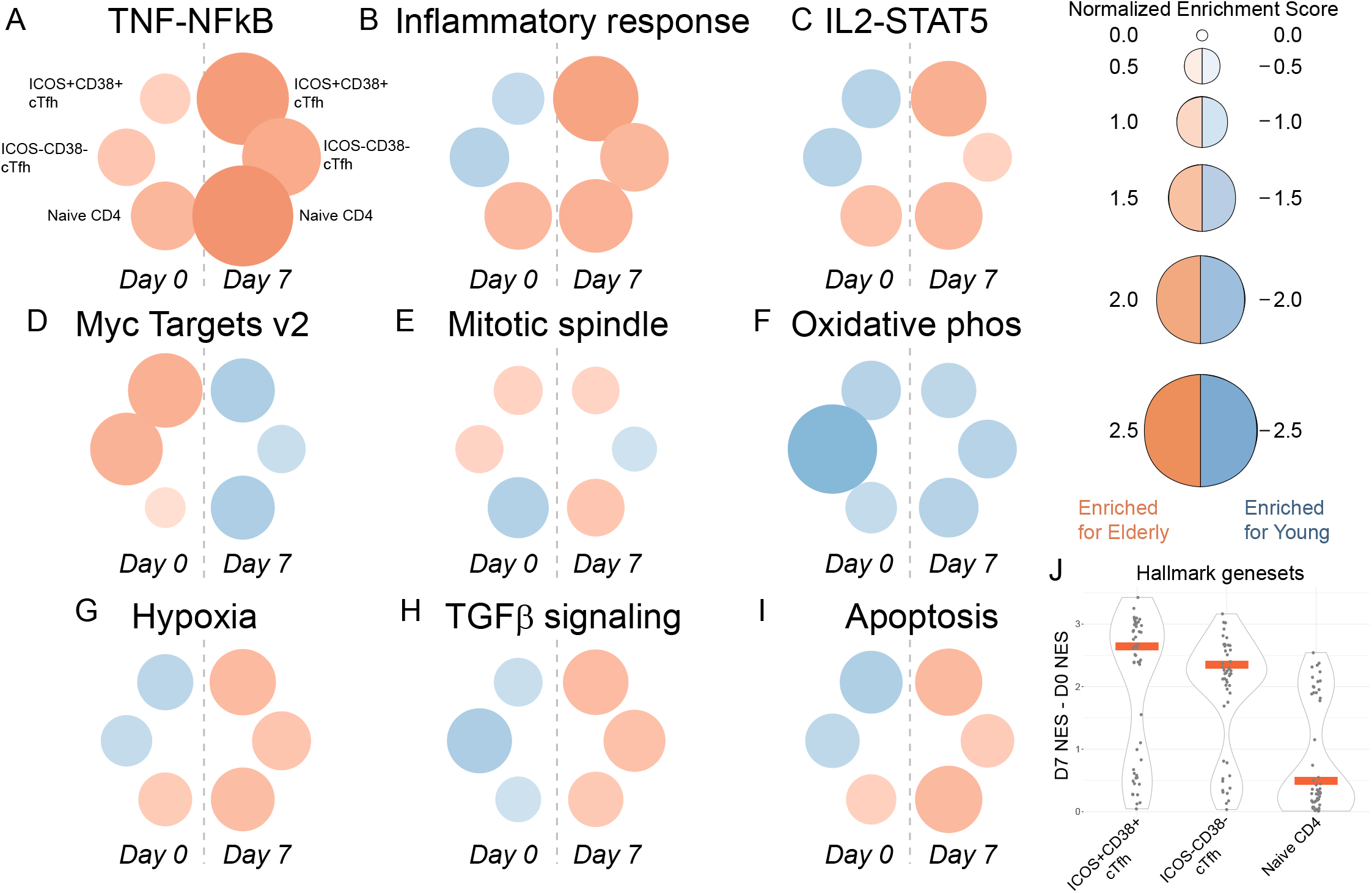
Age-related differences in signaling pathways are preferentially revealed by subset and time point. **A-I.** GSEA analyses were performed to compare young versus elderly for each CD4 T cell subset for each time point for the Hallmark collection in MSigDB. Normalized enrichment score is shown as circle size and color, with redder colors indicating greater enrichment towards elderly and bluer colors indicating greater enrichment towards young. **J.** A “delta-NES” score was calculated by subtracting the day 0 NES score from the day 7 NES score with aging for each Hallmark geneset for each subset. Each dot represents one geneset. Red bar indicates the median for each group.

### The different signatures of ICOS+CD38+ cTfh from young and elderly subjects are most apparent at day 7 post vaccination

Differences in the transcriptional profiles CD4 T cells with aging were most prominent in ICOS+CD38+ cTfh at the cellular pathway level following influenza vaccination (**Figure 5**). We next wanted to determine whether a whole-blood baseline gene signature of aging would be reflected in the same manner and whether vaccination was necessary to identify features of aging.

To test this idea, baseline whole-blood microarray data from the full study cohort (27 young and 35 elderly, Immport SDY739) was used to assess differential gene expression with aging, using the same cohort from which the subjects for the RNAseq (**Figures 1–5**) were derived. From these data, we constructed signatures of youth (354 genes upregulated in the young) or aging (232 genes upregulated in the elderly) (**Supplemental Table 6-7, Supplemental Figure 6A**). These signatures were then used to test whole-blood microarray data from other years’ cohorts from the same study *(48, 49)*. We found strong enrichment of the signatures of youth and aging in the young and elderly adults, respectively (**Figure 6A**). The signature of youth had gene ontology terms for RNA metabolism whereas the signature of aging had terms for fatty acid metabolism and cytoskeletal regulation (**Supplemental Figure 6B**), in agreement with other signatures of aging *(28)*.

**Figure 6.**
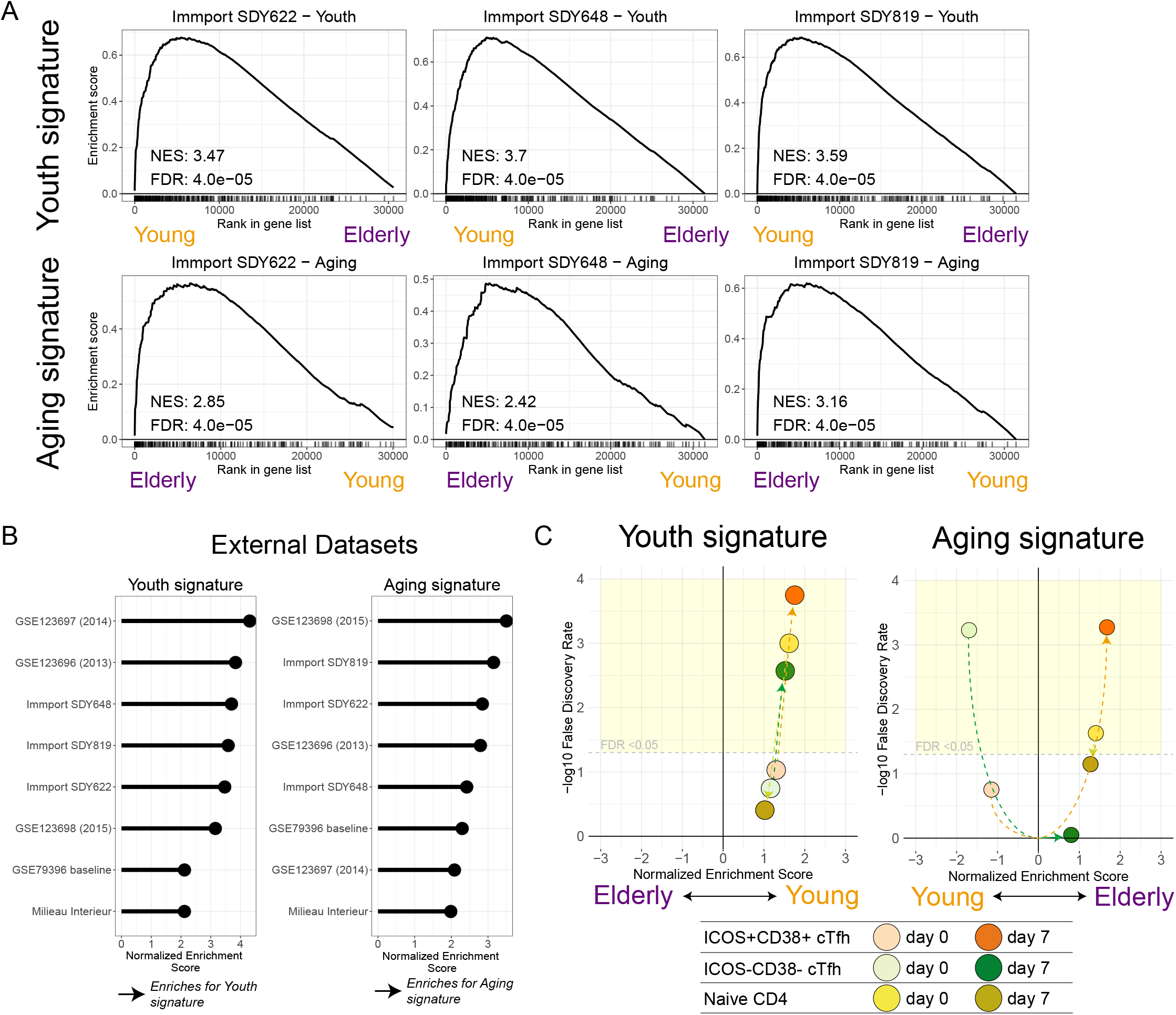
ICOS+CD38+ cTfh transcriptional profiles reveal signatures of aging. **A.** A signature of youth (upper) and aging (lower) was constructed from the differential expression of young versus elderly whole-blood transcriptional profiling from Immport study SDY739. These signatures were tested by pre-ranked GSEA for other vaccine years in this study (SDY622, SDY648, and SDY819). **B.** The signatures of youth and aging were validated against publicly available whole-blood transcriptional profiles (GSE accessions 123696, 123697, 123698, and 79396) and Nanostring target profiling (EGAS00001002460, Milieu Interieur) for young (ages 20-40) and elderly (age 60+). NES are shown for youth (left) and aging (right) signatures. **C.** CD4 T cell subsets were tested for signatures of youth and aging before and after influenza vaccination. Naïve (yellow shades), ICOS-CD38- cTfh (green shades), and ICOS+CD38+ cTfh (orange shade) are shown, with days 0 and 7 connected by dotted lines. Yellow shaded region indicates FDR<0.05 for reference.

To further validate these signatures, we identified several published studies *(13, 50–52)* that generated whole-blood transcriptional profiles on adults under age 40 or over age 65 (**Supplemental Table 8**). From these data, we performed differential expression analysis of young compared to elderly in multiple geographically distinct studies and again found robust enrichment for these signatures in the external datasets (**Figure 6B, Supplemental Figure 6C**). To test these signatures on continuous (rather than cohorted) data, GSVA scores for the signatures of youth and aging were compared against the Nanostring data for subjects in a large cohort of humans (n=986) across lifespan (Milieu Interieur cohort). In this cohort, the signature of youth negatively correlated with chronological age (Pearson r=−0.28, P=8.6×10^−20^, n=986) and the signature of aging positively correlated with chronological age (Pearson r=0.31, P=4.6×10^−23^) (**Supplemental Figure 6D**). These results demonstrate conserved gene expression patterns for youth and aging in whole-blood transcriptional profiling data in multiple published studies.

We next tested whether the signatures of youth and aging could be discerned in the CD4 T cell subsets examined above before and after influenza vaccination. For example, the signature of youth was positively enriched in the young cohort in all subsets at day 0. However, this signature of youth was most strongly enriched in the ICOS+CD38+ cTfh at day 7 after vaccination (**Figure 6C, Supplemental Figure 6E**). Similarly, the signature of aging was evident at day 0, such as in naïve CD4 T cells from the elderly. Again, however, the strongest enrichment of the aging signature was revealed in the ICOS+CD38+ cTfh after vaccination (**Figure 6C, Supplemental Figure 6E**). Together, these data demonstrated that ICOS+CD38+ cTfh are susceptible to immunologic perturbation with influenza vaccination and sensitively reflect general immunological or organismal environment of youth and aging in the cellular transcriptional program.

## Discussion

Induction of protective antibody responses is the goal of most vaccines. Many vaccines are less effective in the elderly, but the underlying causes including how aging impacts Tfh and B cell responses remain poorly understood. In this study, we examined the cTfh response to vaccination in young and elderly adults to interrogate the effects of aging on Tfh. Although ICOS+CD38+ cTfh responses were similar in magnitude in young and older adults, many major cellular pathways were altered and these changes were apparent in the transcriptional signatures and transcriptional networks of these cells. There were several key observations from these studies. First, despite similar magnitude of responses to vaccination, ICOS+CD38+ cTfh from young and elderly subjects differed dramatically in pathway and network level signatures with more prominent regulation of metabolism and cell cycle in cTfh from young subjects and an increase in signatures of inflammation and cytokine signaling in the elderly. Second, signatures of inflammatory responses, IL2-STAT5 signaling and TNF-NFkB signaling in particular were substantially stronger in ICOS+CD38+ cTfh from elderly subjects. Third, vaccine-induced ICOS+CD38+ cTfh reflected underlying signatures of aging versus youth more strongly than ICOS-CD38- cTfh or naïve CD4 T cells suggesting that recently activated cTfh may function as biosensors of underlying inflammatory and/or host physiological environments and reflect underlying features of “immune health”.

Increased baseline inflammation in the elderly has often been associated with declines in overall health *(53)* and, in particular, the TNF-NFkB pathway has been implicated in many pathological changes with aging. However, these correlations are at odds with the apparent detrimental effects of TNF pathway blockade on vaccination *(20–22)* that suggest that blocking residual TNF signaling impairs, rather than enhances, vaccine-induced antibody responses. Moreover, data from mice indicates a key role for TNF in GC responses *(23–26)*. Thus, how TNF-NFkB signaling regulates GC-dependent, vaccine-induced immunity in the elderly remains incompletely understood. Here, TNF-NFkB signaling was correlated with quantitatively better antibody responses suggesting a positive role for TNF signaling in human GC-dependent immune responses. ICOS+CD38+ cTfh displayed a strong transcriptional signature of TNF-NFkB pathway signaling in all subjects, especially the elderly, and blockade of TNF signaling *in vitro* led to reduced immunoglobulin production by B cells *in vitro*. Furthermore, these *in vitro* studies indicated reduced survival of cells upon blockade of TNF signaling. Indeed, our results point to a pro-survival effect of TNF signaling likely due to effects of TNF signaling on cTfh and/or TNF production by cTfh. Despite increased expression of pro-apoptotic Bcl family members such as *BID* in ICOS+CD38+ cTfh, there was a concomitant increase in expression of pro-survival factors *BCL2A1* and *MCL1* that strongly correlated with TNF-NFkB GSVA scores in ICOS+CD38+ cTfh at day 7 suggesting a TNF-NFkB-dependent survival signal in activated cTfh. These genes have NF-kB binding sites in their respective promoter sequences *(54, 55)* linking TNF signaling to a potential pro-survival circuit in these cells. Moreover, TNFR1 expression in ICOS+CD38+ cTfh at day 7 positively correlated with neutralizing antibody production. These data are in agreement with, and may provide an explanation for, clinical data from patients receiving therapeutic TNF blockade demonstrating fewer GC reactions *(56)* and attenuated antibody responses to influenza vaccination (22). These results may have direct implications for older adults receiving therapeutic TNF blockade, who may thus be at even greater risk for vaccine non-responsiveness. Although these data suggested that the defects in vaccination in the elderly cannot be ascribed to TNF-NFkB activity in Tfh leading to reduced Tfh activity, it is possible that excessive Tfh activity impairs effective selection of high-quality antibody producing B cell clones. These results may also have implications for increases in autoreactive antibodies with age and/or age associated autoimmune diseases.

Although many circulating immune cell types can be altered with aging, our focus here on cTfh and CD4 T cells revealed a novel feature of vaccine-induced cTfh. By focusing on the ICOS+CD38+ cTfh population on day 7 after influenza vaccination, we found that these cells preferentially reflected underlying inflammatory and age-related biological signatures compared to ICOS-CD38- cTfh or naïve CD4 T cells. There are several potential reasons for this feature of ICOS+CD38+ cTfh. For example, the increase in the ICOS+CD38+ cTfh subset at day 7 post vaccination likely reflects an induction and/or reactivation of influenza-specific CD4 T cells, as this subset is enriched in influenza antigen-specific cells and similar TCR clones are recalled into this population upon yearly vaccination *(35)*. Thus, it is possible that these influenza specific T cells are qualitatively different from the ICOS+CD38+ cTfh circulating at baseline. However, a second related feature of these vaccine-induced cTfh relevant to these signature enrichments may be the synchronous nature of the responses. The ICOS+CD38+ cTfh responses at baseline likely reflect either ongoing chronically activated cTfh or cTfh in different phases of acute activation. In contrast, the vaccine-induced cTfh are aligned in their activation and differentiation kinetics, perhaps resulting in more focused transcriptional circuitry. Recent activation might also facilitate sensitivity and/or changes in inflammatory cytokine signaling as has been suggested for NFkB signaling in GC B cells *(57)*. In addition to enrichment for TNF-NFkB signaling and a broader signature of aging, activated cTfh also displayed transcriptional evidence of age-dependent alterations in many other pathways including other inflammatory circuits like IL2-STAT5 signaling, TGFβ pathway signaling, IL6-JAK-STAT3 signaling and several non-immune pathways. These observations were notable given the importance of both IL-2 and TNF-NFkB in Tfh biology and in inflammation *(58–61)*. To aid in our understanding of the impacts of aging on cTfh, we constructed transcriptional signature maps of youth and aging based on whole-blood transcriptional profiling from 27 young and 34 elderly adults. These signatures were then validated using multiple geographically distinct cohorts, representing a total of 566 young adult samples and 344 elderly adult samples. The GSVA scores for the signatures of youth and aging from these dichotomous studies correlated with chronological age as a continuous variable in the Milieu Interieur study of nearly 1000 adults, with chronological age predicting 7.8-9.6% of the variance in the GSVA scores of our signatures. These analyses identified differential expression of RNA metabolism and regulation of cellular migration in the young and elderly, respectively, similar to other studies that have established signatures of aging *(28)*. Overall, these data indicated that the highest enrichment scores for our signatures of youth and aging were in the day 7 ICOS+CD38+ cTfh subset, demonstrating the advantage of profiling this subset to interrogate aging in the context of vaccine perturbation. These observations suggest that cTfh, and especially the synchronized, vaccine-induced cTfh at day 7 after influenza vaccination, may be sensitive biosensors of underlying features of immune inflammation or immune health.

Studies of the effects of aging on the vaccine-induced immune response are needed to understand poor immune responses in aging adults who may be at heightened risk from infection. Rational vaccine strategies will require knowledge of how alterations in background host physiology, such as aging, impacts coordination of germinal center dependent antibody development and humoral immunity. These studies not only identify novel roles for inflammatory pathways in alterations of cTfh during aging, but also point to vaccine induced cTfh as a cell type that might provide a unique window into changes in overall immune health and fitness.

## Materials and Methods

### Human subjects

In the Fall of 2014, study subjects were recruited and consented at the Clinical Research Unit at Duke University Medical Center (Durham, NC, USA), in accordance with the Institutional Review Boards of both Duke University and the University of Pennsylvania (Philadelphia, PA, USA). Subjects were classified as young (30-40 years of age) or elderly (65 years of age or older). Subjects were eligible if they were community-dwelling and had not received influenza vaccine in the prior 6 months; they were excluded if they had contraindications to influenza vaccine, active substance abuse, HIV/AIDS, clinically active malignancy, immunomodulatory medication need (i.e. chemotherapy, corticosteroids), or active illness (i.e. active respiratory tract infections). Seasonal trivalent influenza vaccine (Fluvarix, GlaxoSmithKline) was administered and peripheral venous blood was drawn on days 0, 7, and 28 after vaccination. Blood was collected into heparinized tubes and shipped overnight to Philadelphia, PA. Samples used in **Figure 3A** were from subjects who provided informed consent in accordance with the Institutional Review Boards of the Louis Stokes Veterans Affairs Medical Center and Case Western Reserve University, as previously reported *(32)*.

### Flow cytometry

PBMC and plasma were isolated using Ficoll-Paque PLUS (GE Healthcare) and stained for surface and intracellular markers. Permeabilization was performed using the Intracellular Fixation/Permeabilization Concentrate and Diluent kit (ThermoFisher). Antibodies and clones are described in **Supplemental Table 2**. Cells were resuspended in 1% para-formaldehyde until acquisition on a BD Biosciences LSR II or a BD Symphony A5 cytometer. Fluorescence-minus-one controls were performed in pilot studies. DiOC6 (Fisher) staining was performed according to manufacturer instructions.

### Co-culture experiments

For each co-culture condition, 4.5×10^4^ sorted T cells were combined with 5×10^4^ naïve B cells (defined as CD3^−^CD19^+^CD27^lo^IgD^hi^). Co-cultures were either left unstimulated, stimulated with Staphyloccocal Enterotoxin B (SEB, 0.5 μg/mL), SEB (0.5 μg/mL) with human TNF cytokine (Biolegend, 125 ng/mL), or SEB (0.5 μg/mL) with αTNF antibodies (BD Biosciences, clone MAb11, 2 μg/mL). At the conclusion of the co-culture, supernatant was aspirated and saved. The remaining cells underwent anti-human Fc-blockade (Biolegend, TruStain FcX) for 10 minutes at room temperature, followed by routine staining for flow cytometry. Cultures with B cells only were stimulated using soluble αIgM (10 ug/mL, clone DA4-4, Fisher) with soluble trimeric MegaCD40L protein (100 ng/mL, Enzo Life Sciences).

### Multiplex bead assays

Multiplex bead assays for plasma samples were performed in duplicate for TNF from the MILLIPLEX MAP Human Cytokine/Chemokine Magnetic Bead Panel (Millipore Sigma). Co-culture supernatants were assayed in duplicate for IgG1 production using the Legendplex Human Immunoglobulin Isotyping Panel (Biolegend) and data acquired on a BD Symphony A5 cytometer, in accordance with manufacturer instructions.

### Transcriptomic analyses

PBMC were sorted on a BD Aria II cell sorter, followed by total RNA extraction by RNeasy Micro Plus kit (Qiagen) and polyA amplification with SMARTer Ultra-Low Input RNA kit v3 (Clontech) according to manufacturer instructions. Libraries were prepared using the Illumina Nextera XT Library Preparation kit and sequenced on an Illumina HiSeq 2000 using 100bp single-read format. FASTQ files were trimmed with Trimmomatic (version 0.32), aligned using STAR (version 2.5.2a) *(62)* and normalized by PORT (https://github.com/itmat/Normalization/, version 0.8.5) against the GRCh38 reference assembly of the human genome. Variance-stabilizing transformation and differential expression were performed using DESeq2 (version 1.20.0) *(63)* based on genes with at least 20 counts in at least 25% of all libraries, using the R environment (version 3.5.0). Of the 84 resultant libraries, one sample was excluded from further analyses (**Supplemental Figure 1F-G**). Plots were made by ggplot2 (version 2.2.1) and heatmaps using the inferno color-scheme from the “viridis” library. For differential expression analysis, genes were included if they had at least 20 counts in at least 25% of the samples. Samples from the same subject (e.g. inter-subset comparisons or day 0-to-day 7 comparisons) were treated as paired in statistical models for DESeq2. t-SNE maps were produced using Rtsne (version 0.13) from the transformed counts data. Gene set enrichment analyses *(37)* were performed with at least 1000 permutations of pre-ranked GSEA (http://software.broadinstitute.org/gsea/downloads.jsp). Gene ontology was performed in Cytoscape using ClueGo *(64, 65)* or with Metascape (https://metascape.com) *(66)*. Ingenuity Pathway Analysis (Qiagen) was performed by uploading tables of the differential expression results from DESeq2 analyses for log_2_ fold-change, p-values, and p-adjusted values, followed by Expression Analysis core analysis for all genes with p-value < 0.10.

### Network analysis

Weighted gene correlation network analysis (WGCNA, version 1.63) *(40)* was performed on variance-stabilizing transformation data, as calculated by DESeq2 *(63)* after filtering out genes with less than 5 counts in fewer than 12% of all libraries. Adjacency matrix was calculated using β=20. Minimum module size was 100. Gene ontology was analyzed using Metascape *(66)* for all genes with a module membership of at least 0.8. Pre-ranked GSEA was performed using all genes ranked by membership for each module. Transcription factors were identified from a curated list *(41)* and displayed with functional association data overlaid from GeneMANIA *(67)*.

### Aging signature

Transcriptional profiling data for whole blood microarray studies was obtained from the Gene Expression Omnibus (GEO accession numbers 123687, 123696, and 123698), from Immport (SDY622, SDY648, SDY739, and SDY819), and from the European Genome-phenome Archive (EGAS00001002460). For each dataset, subjects were stratified into cohorts for young adults (<40 years old) or elderly adults (>65 years old). Differential expression was calculated using GEO2R or by limma-voom *(68)*. An aging signature of 354 genes was constructed from genes downregulated in the elderly cohort from SDY739 with p.adj <0.20 using limma-voom. For analyses of the aging signature in CD4 subsets, pre-ranked GSEA analyses were performed with at least 25000 permutations and in subsets where FDR result was 0, the FDR was instead reported as 1/permutations or 4×10^−5^ as per the GSEA Wiki FAQ entry 3.6.

### Influenza-specific antibodies

The two influenza A vaccine strains of the 2014/2015 seasonal influenza vaccine, A/California/7/2009 (H1N1) pdm09-like virus and A/Victoria/361/2011 (H3N2)-like virus, were obtained from the Centers for Disease Control and Prevention (Atlanta, GA). Assays to detect hemagglutinin inhibition assay (HAI) titers and microneutralization (MN) titers were performed. Infectious virus was used for neutralizing Ab assays or inactivated by β-propionolactone for H1N1/California- and H3N2/Victoria-specific binding Ab ELISA assays. Nunc Maxisorp plates (Nunc) were coated with 10 mg/mL influenza A/H1N1/California and A/H3N2/Victoria virus along with isotype standards for IgA1, IgG, and IgM (Athens Research and Technology) in bicarbonate buffer overnight at 4°C. Plates were blocked with 3% BSA in PBS and incubated with heat-inactivated sera of young and elderly subjects. Abs were detected using alkaline phosphatase-conjugated mouse anti-human IgA1, IgG, and IgM (Southern Biotechnology).

### Statistics

Statistical analyses were performed with Prism 7 (GraphPad) and R. Data was compared using Student’s *t* test, paired *t* test, one-way Analysis of Variance (ANOVA) with Tukey post-hoc analysis, non-parametric ANOVA with Friedman’s posttest, or Fisher’s Exact test, as indicated. All t-tests were performed as two-tailed tests at a 0.05 significance level. Outlier analysis for **Figure 1E** was performed using Grubb’s test at α=0.05 and the plot with all points included is shown in **Supplemental Figure 1E**.

### Data and code

Transcriptional profiling data reported here are available at the Gene Expression Omnibus (GEO) under accession number GSE 134416. R scripts used in analyses and figure generation are available on Github (https://github.com/Sedmic/cTfh_AgingSignature).

## Acknowledgements

We would like to thank the subjects who participated in our studies, as well as members of the Wherry lab who provided critical feedback in this project. We would also like to thank the Next Generation Sequencing Core, the Flow Cytometry Core, and the Human Immunology Core at the University of Pennsylvania for their advice and assistance in these experiments. This work was supported by NIH grants AI114852 and AG047773 to R.S.H, the Pediatric Infectious Diseases Society Fellowship Award, the National Center for Advancing Translational Sciences of the National Institutes of Health under award number KL2TR001879 to L.A.V., and a Mentored Research Scholar Award from the Penn Center for AIDS Research (CFAR) to L.A.V., an NIH-funded program (P30 AI 045008), as well as R50 CA211199 (to A.V.K.). K.E.S. was supported in part by the NIH/National Institute on Aging Claude D. Pepper Older Americans Independence Centers (grant AG028716). D.H.C. was supported by the Veterans Affairs and AI108972. This work was additionally supported by NIH grants AI105343, AI112521, AI082630, AI201085, and AI117950 to E.J.W. as well as U.S. Broad Agency Announcement HHSN272201100018C (to K.E.S., H.C.J.E., and E.J.W.). E.J.W. is also supported by the Parker Institute for Cancer Immunotherapy which supports the cancer immunology program at UPenn.

## Author contributions

R.S.H., L.V.S., L.A.V., A.M., B.B., J.K., and D.H.C designed and performed experiments. R.S.H., and E.J.W. conceived the overall design. S.K., R.K., and H.E. performed and analyzed antibody assays. A.K. analyzed Immport studies SDY622, SDY648, SDY739, and SDY819. C.A. assisted with analysis of the Milieu Interieur data. S.D. and K.E.S. recruited and vaccinated study subjects. All authors analyzed and interpreted data, discussed the results, and commented on the manuscript. R.S.H., and E.J.W. wrote the manuscript.

