## Supplemental Figures for "Vaccine-induced ICOS+CD38+ cTfh are sensitive biosensors of age-related changes in inflammatory pathways"

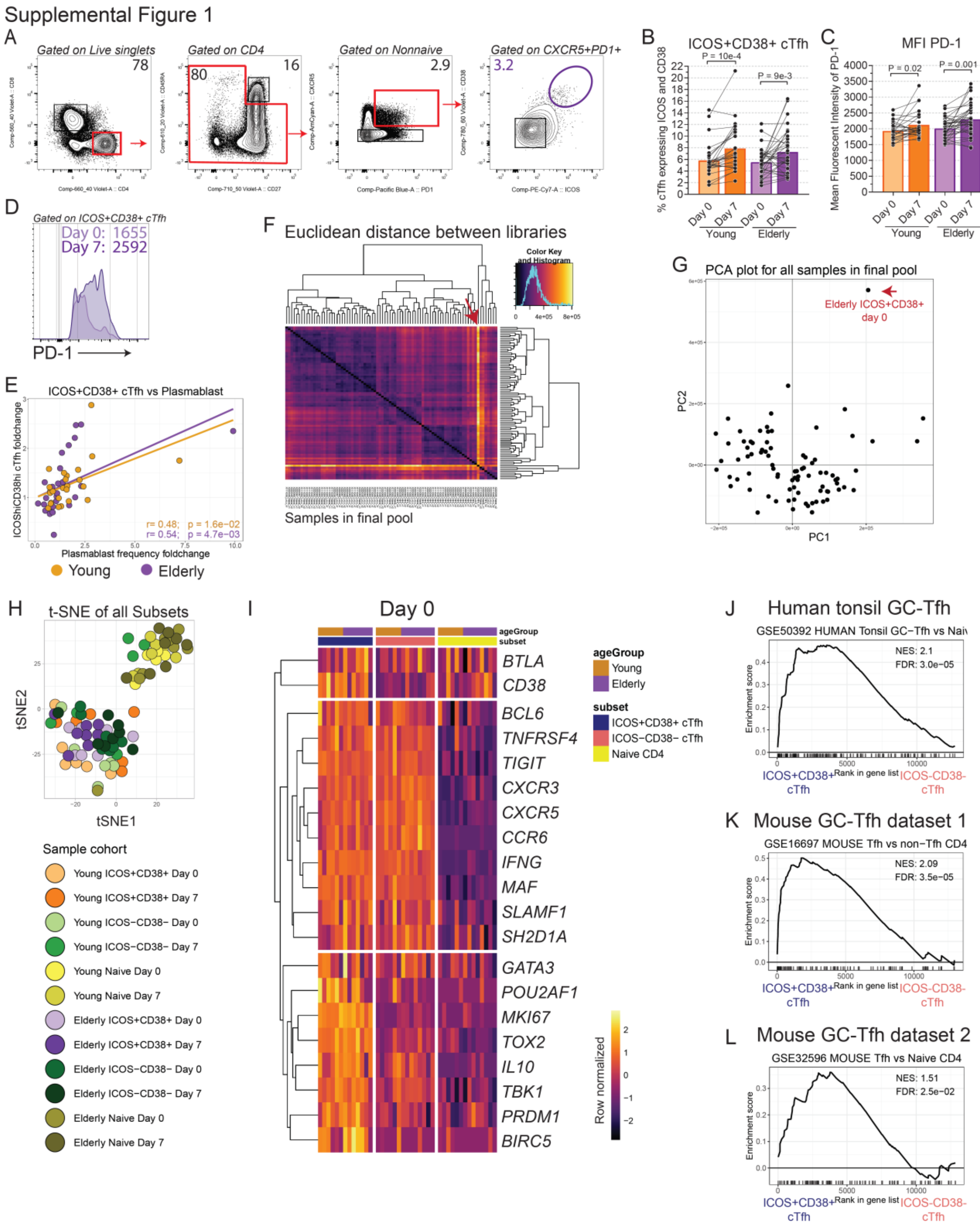

**Supplemental Figure 1. A.** Flow cytometry gating scheme shown for identifying cTfh subsets from peripheral blood. **B.** Summary plots shown for frequency of cTfh co-expressing ICOS and CD38 for young (orange, $P=10^{-3}$, paired t-test, n=27) and elderly (purple, $P=8.6x10^{-3}$, paired t-test, n=35) at days 0 and 7 after vaccination. **C.** Summary plots shown for Mean Fluorescence Intensity of PD-1 (CD279) in ICOS+CD38+ cTfh for young (orange, P=0.015, paired t-test; n=27) and elderly (purple, $P=10^{-3}$; paired t-test; n=35) at days 0 and 7 after vaccination. **D.** Example PD-1 stain shown for one subject at day 0 (light purple) and 7 (dark purple) after influenza vaccination. Plot gated on ICOS+CD38+ cTfh. Geometric mean fluorescent intensity shown. **E.** Correlation shown for the ICOS+CD38+ cTfh frequency fold-change from day 7 compared to day 0, vs the plasmablast response fold-change in frequency from day 7 compared to day 0. Shown is the full dataset with all outliers included for young (orange) and elderly (purple) subjects. **F.** Euclidean distance matrix calculated for all samples in the final pool for RNAseq after raw data processing. One outlier was identified by the red arrow. **G.** Principal component analysis on all samples in the final pool for RNAseq after raw data processing. The outlier (same as in **F**) is indicated by arrow and was excluded from all subsequent analyses. **H.** t-Stochastic neighbor embedding (t-SNE) plot of remaining 83 samples. **I.** Log-transformed transcriptional profiling data was queried for selected genes for young (orange bars) and elderly (purple bars) for CD4 subsets at day 0 after vaccination. Each column represents one unique subject. Heatmap is row-normalized. Row gaps and column gaps are based on hierarchical clustering and CD4 subsets, respectively. **J-L.** Normalized Enrichment Score (NES) and False Discovery Rate (FDR) q-value are shown for the pre-ranked gene set enrichment analysis (GSEA) comparison of ICOS+CD38+ cTfh to ICOS-CD38- cTfh at day 0 for all subjects combined, for GSE50392 (human tonsillar Tfh vs naïve CD4) (**J**), GSE16697 (mouse Tfh vs non-Tfh CD4) (**K**), and GSE32596 (mouse Tfh vs naïve CD4) (**L**).

Supplemental Figure 2

A

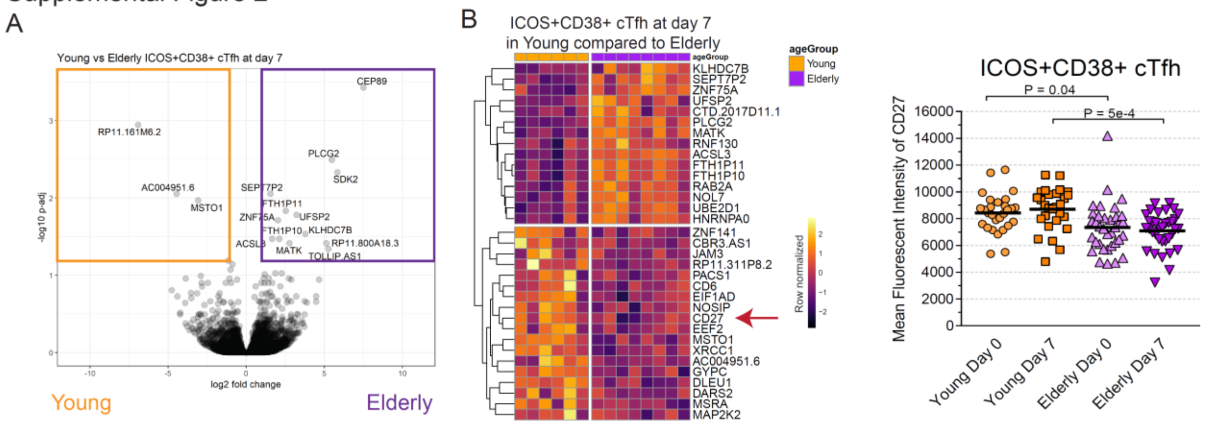


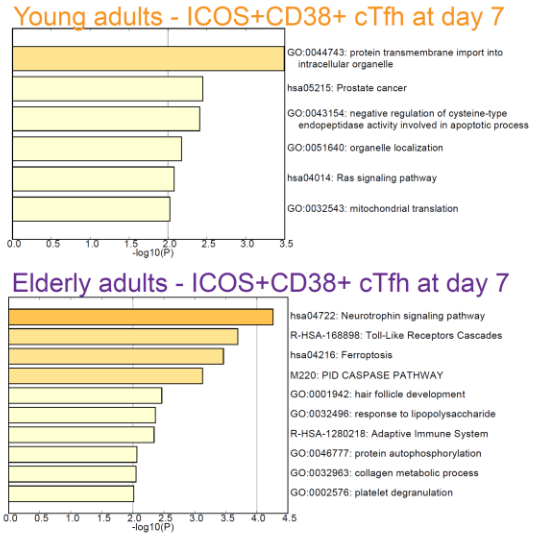


C

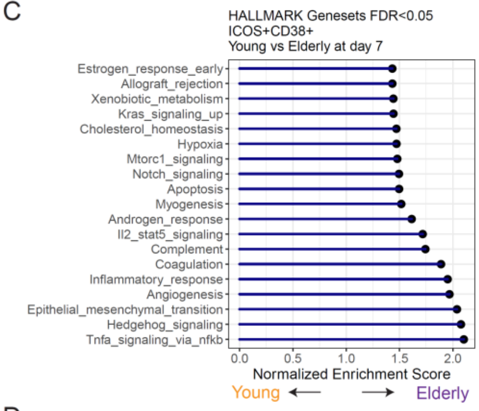


E

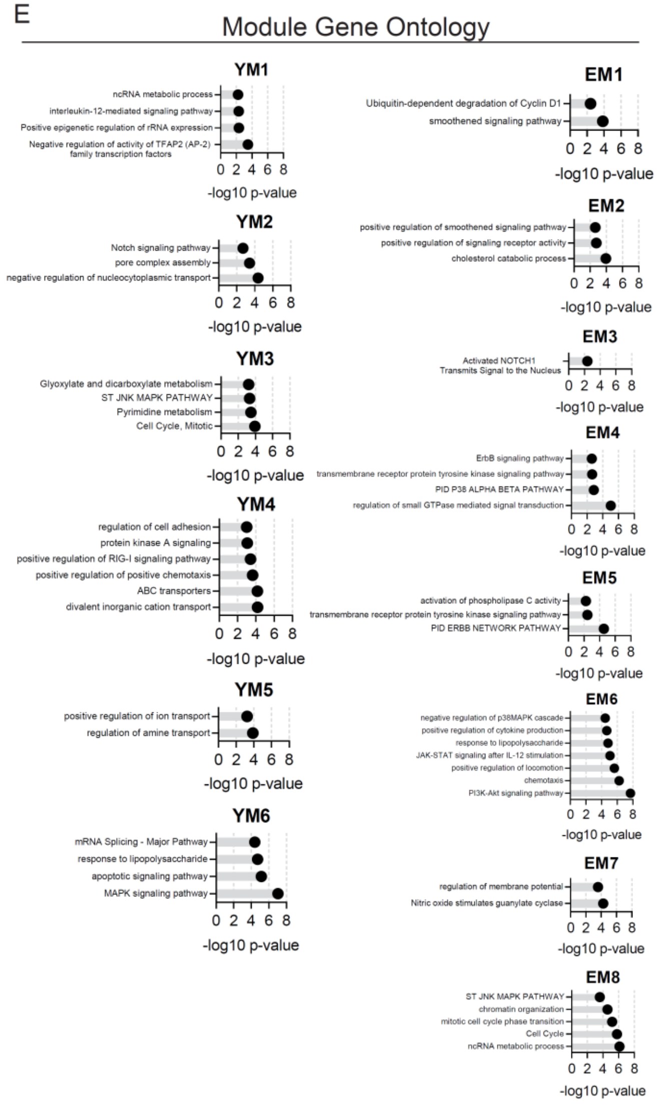


D

Young adults

| Module | Label | # Genes |
|---|---|---|
| Green | YM1 | 655 |
| Red | YM2 | 424 |
| Brown | YM3 | 790 |
| Turquoise | YM4 | 1610 |
| Yellow | YM5 | 690 |
| Blue | YM6 | 924 |

Elderly adults

| Module | Label | # Genes |
|---|---|---|
| Light green | EM1 | 426 |
| Green | EM2 | 813 |
| Brown | EM3 | 1155 |
| Pink | EM4 | 629 |
| Black | EM5 | 647 |
| Turquoise | EM6 | 1547 |
| Yellow | EM7 | 890 |
| Blue | EM8 | 1503 |

Supplemental Figure 2

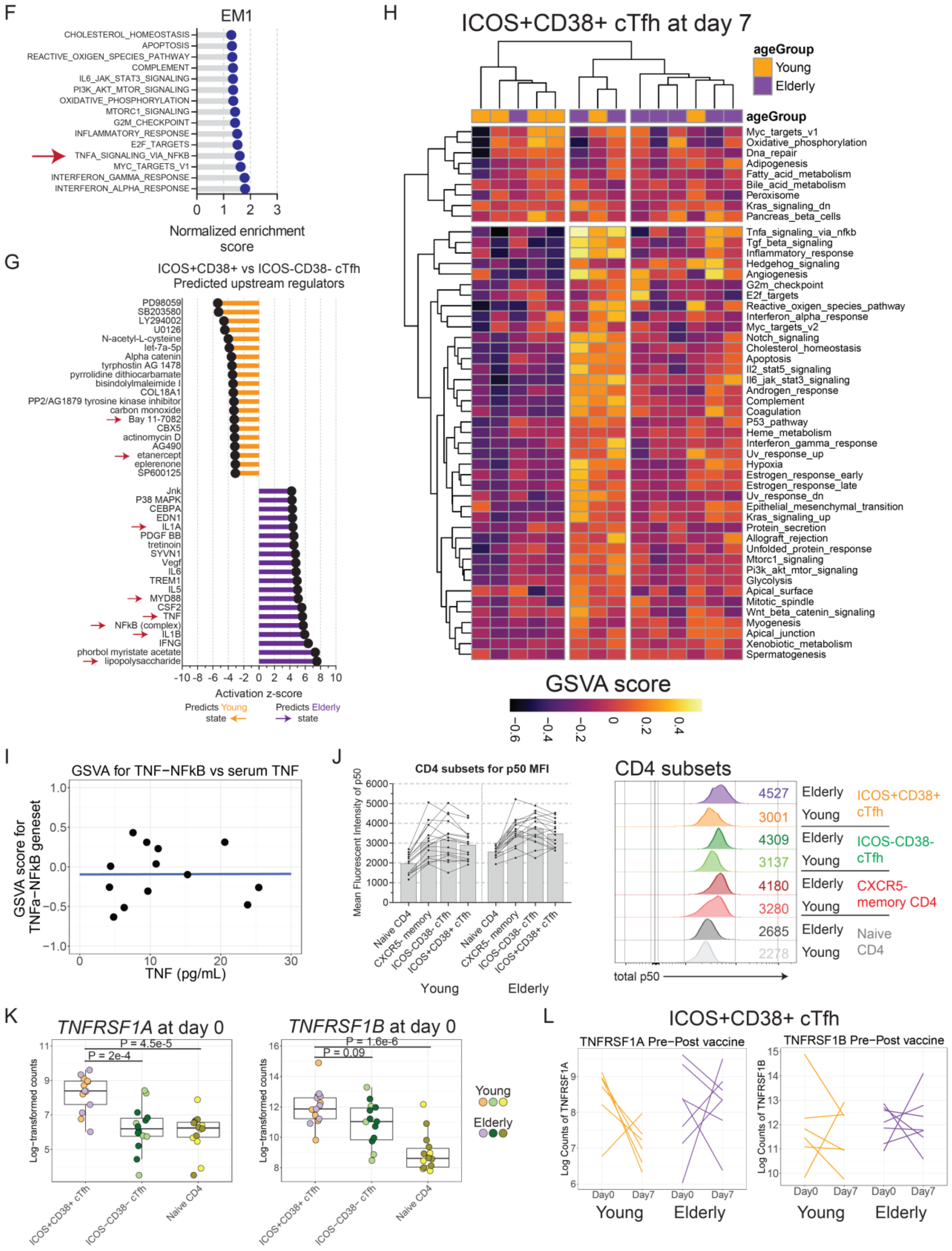

**Supplemental Figure 2. A.** Volcano plot (top) shown for the differential expression analysis of ICOS+CD38+ cTfh from young adults and elderly adults at day 7 after vaccination. Gene ontology was performed on the 100 most differentially-expressed genes for young (middle) or elderly (bottom) subjects. **B.** Log-transformed transcriptional profiling data (left) was queried for differentially-expressed genes for the ICOS+CD38+ cTfh at day 7 from young (orange) and elderly (purple). Each column represents one unique subject. Heatmap is row-normalized and gaps are based on hierarchical clustering. Geometric mean fluorescence intensity of CD27 (right) is shown for the ICOS+CD38+ cTfh at days 0 and 7 (one-way ANOVA with Tukey's post-test; n=28 for young, n=35 for elderly). **C.** Pre-ranked GSEA was used to compare ICOS+CD38+ cTfh at day 7 from young vs elderly subjects. Pathways with FDR <0.05 are shown. Positive NES indicate greater enrichment in the elderly than young. **D.** Weighted gene correlation network analysis was performed for ICOS+CD38+ cTfh at day 7 after vaccination for young adults and elderly adults separately. The six young adult modules and eight elderly adult modules were relabeled as indicated. **E.** Gene ontology by Metascape was performed for all genes in each module with module membership >0.80. **F.** Pre-ranked GSEA was performed on module EM1 using the Hallmark gene sets from MSigDB. TNF-NFkB geneset is indicated by the red arrow. **G.** Ingenuity Pathway Analysis was used to assess predicted upstream regulators in the comparison of ICOS+CD38+ cTfh in young and elderly subjects at day 7 after vaccination. Top 20 highest and 20 lowest-scoring terms by activation z-score are shown. Arrows indicate terms directly relevant to NFkB signaling. **H.** Gene set variation analysis (GSVA) was performed for ICOS+CD38+ cTfh at day 7 from all 14 subjects for the Hallmark collection in MSigDB. **I**. Scatter-plot shows GSVA score per subject compared to the plasma TNF concentration at baseline. **J.** PBMC were assayed by flow cytometry for total NFkB p50 protein expression. Numbers on histogram indicate mean fluorescence intensity for p50. Summary data (left) and example plot (right) are shown for one young and one elderly subject for p50 protein in the different CD4 subsets. **K.** $Log_2$-transformed mRNA counts shown for gene expression for *TNFRSF1A* (left) or *TNFRSF1B* (right) in CD4 subsets for young and elderly subjects (one-way ANOVA with Tukey's, n=6 or 8). **L.** Gene expression of *TNFRSF1A* (left) and *TNFRSF1B* (right) were assessed from RNAseq data, shown as before-and-after plots for normalized mRNA counts. Each line indicates one unique subject.

Supplemental Figure 3

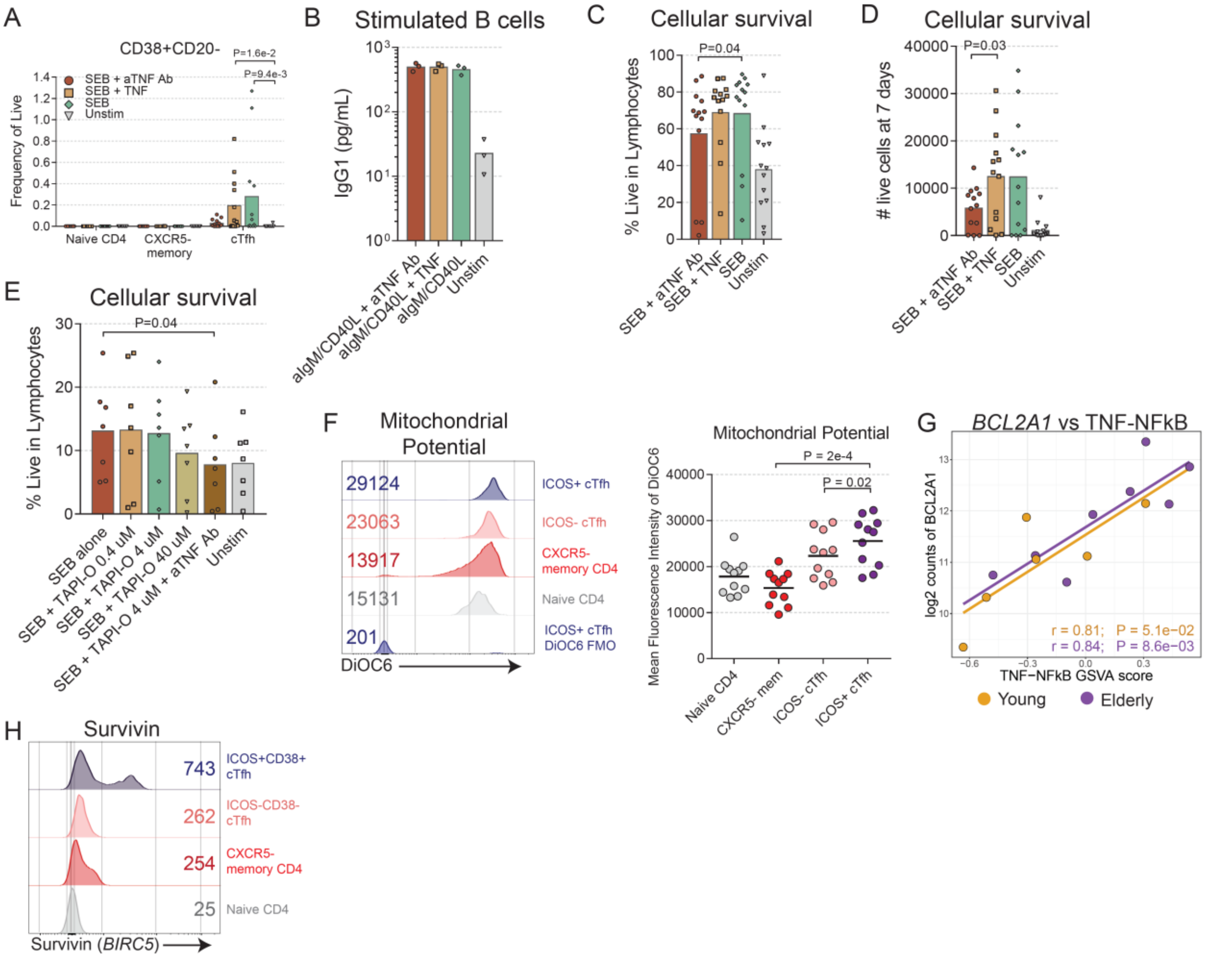


**Supplemental Figure 3. A.** PBMC from young adults were freshly isolated and sorted for co-culture with autologous naïve B cells ($CD3^{-}CD19^{+}CD27^{lo}IgD^{+}$). Plasmablast frequency (gated as $CD19^{+}CD38^{+}CD20^{-}$) as a proportion of live cells was measured after 7 days, as shown for the following conditions: unstimulated (grey), SEB alone (0.5 µg/mL, green), SEB with recombinant human TNF (125 ng/mL, tan), or SEB with αTNF antibodies (2 µg/mL, sienna) (one-way ANOVA with Freidman's test; n=12 per group). **B.** Supernatants from *in vitro* culture of naïve B cells stimulated with αIgM (10 µg/mL) and CD40L trimer (100 ng/mL) with TNF modulation were measured for IgG1 after 7 days. **C-D.** Live cell counts were measured by flow cytometry after 7-day culture of cTfh with naïve B cells as in (**A**) (one-way ANOVA with Freidman's test; n=12 per group), shown as %Live (**C**) and absolute number (**D**). **E.** Coculture was performed with cTfh and autologous naïve B cells stimulated with SEB alone (0.5 μg/mL), or SEB with TAPI-O at the indicated concentrations, or SEB + TAPI-O + αTNF antibodies (2 μg/mL) (one-way ANOVA; n=7 per group), and cells at day 7 are shown as %Live. **F.** Mitochondrial potential was assessed by staining PBMC from young and elderly adults with DiOC6. Example flow cytometry plot (left) and summary data (right) are shown (one-way repeated-measures ANOVA with Tukey's; n=11 per group). **G.** Gene expression of *BCL2A1* from transcriptional profiling of ICOS+CD38+ cTfh at day 7 was correlated against the GSVA scores for TNF-NFkB for young (orange, $P=5.1x10^{-2}$, Pearson r=0.81, n=6) and elderly (purple, $P=8.6x10^{-3}$, Pearson r=0.84, n=8). **H.** Flow cytometry shown for one young subject for expression of survivin protein. MFI is shown.

Supplemental Figure 4

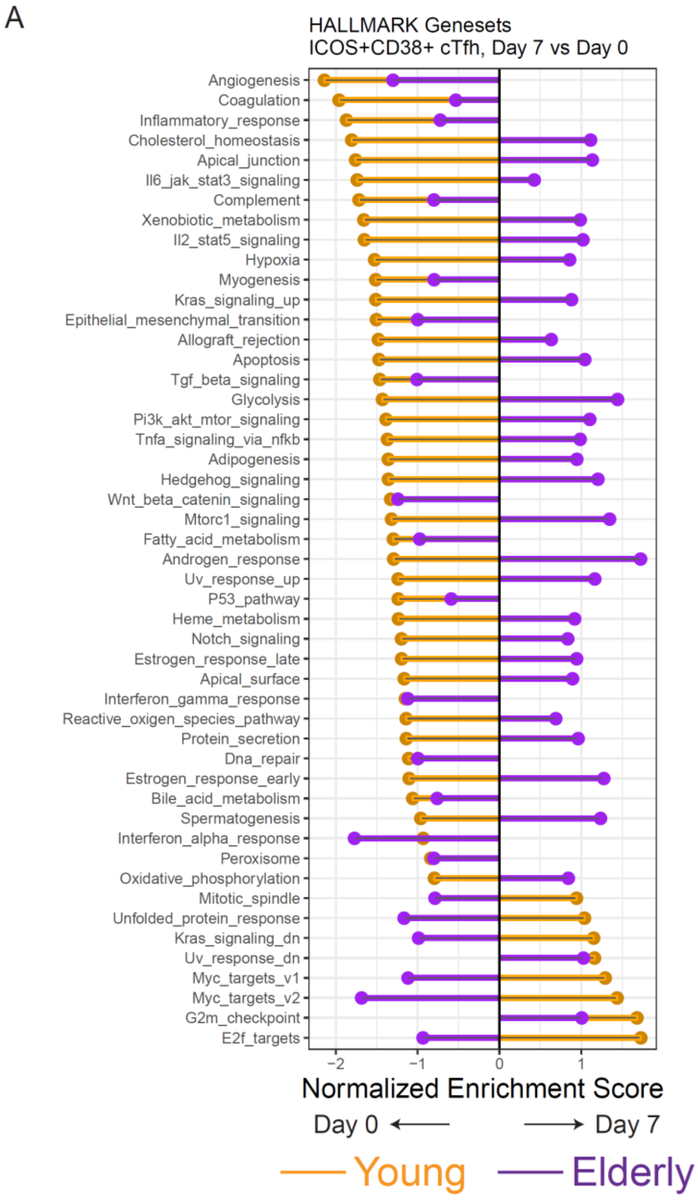


**Supplemental Figure 4.** **A.** Aggregated GSEA results are shown for the full Hallmark genesets collection for the comparison of day 7 vs day 0 for ICOS+CD38+ cTfh in young (orange) and elderly (purple). Positive enrichment scores indicate enrichment at day 7.

Supplemental Figure 5

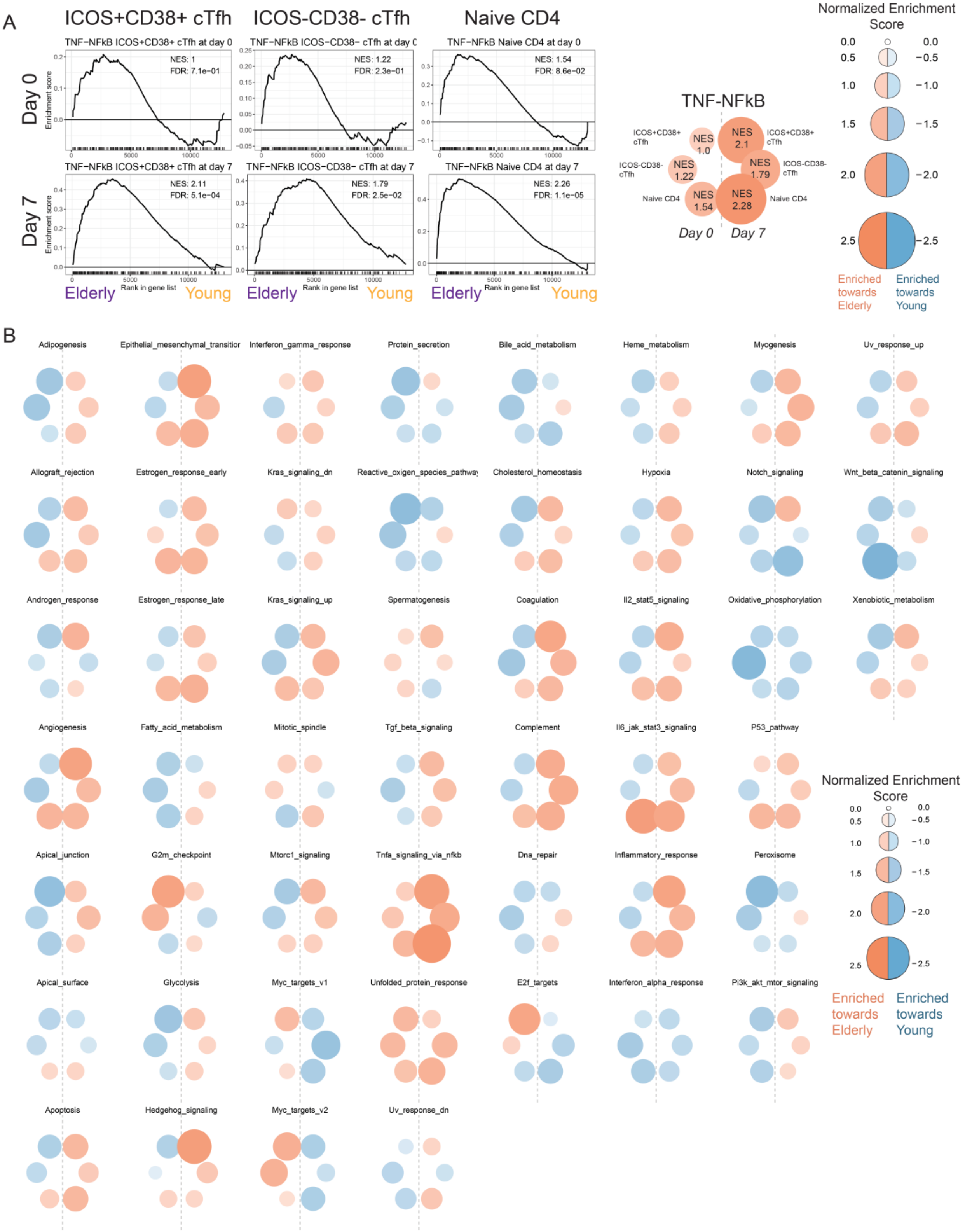

**Supplemental Figure 5.** **A**. Pre-ranked GSEA analyses comparing young and elderly subjects per CD4 subset per time point are shown for the TNF-NFkB gene set. Aggregate hexagon plot for the normalized enrichment scores is shown at right as a prototypical example. **B.** Hexagon plots are shown for the pre-ranked GSEA analyses for all of the MSigDB Hallmark genesets comparing young versus elderly by CD4 subset and by time point.

Supplemental Figure 6

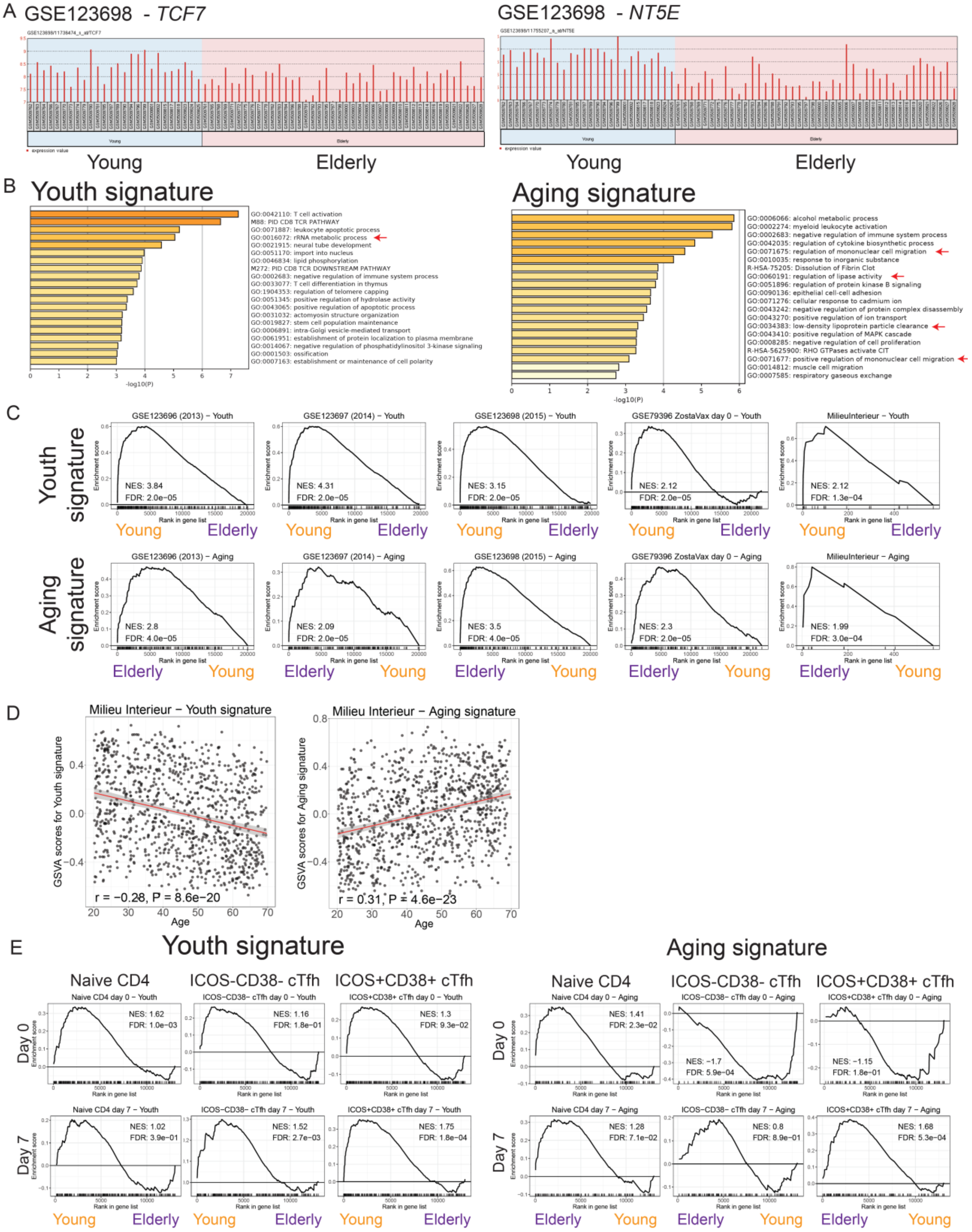

**Supplemental Figure 6.** **A.** Examples shown from GEO2R for GSE123698 for gene expression of *TCF7* (left) and *NT5E* (right) by cohort. **B.** Metascape gene ontology for the youth signature (left) and the aging signature (right). **C.** The youth signature (upper row) and the aging signature (lower row) were tested by pre-ranked GSEA for transcriptional profiling data for the following studies: GSE123696, GSE123697, GSE123698, GSE79396 (day 0 data), and EGAS00001002460 (Milieu Interieur). **D.** The full Milieu Interieur Nanostring dataset was used to test the GSVA scores for the youth and aging signatures against chronological age. The linear regression line (red) is shown for the youth signature (left, $P=8.6\times10^{-20}$, Pearson r=-0.28, n=986) and aging signature (right, $P=4.6\times10^{-23}$, Pearson r=0.31, n=986). **E.** The youth (left) and aging (right) signatures were used to probe CD4 subsets at days 0 and 7 after influenza vaccination by pre-ranked GSEA. NES and FDR are shown.
